# Understanding Language Model Scaling on Protein Fitness Prediction

**DOI:** 10.1101/2025.04.25.650688

**Authors:** Chao Hou, Di Liu, Aziz Zafar, Yufeng Shen

**Affiliations:** Department of Systems Biology, Columbia University Irving Medical Center, New York, NY 10032; Department of Biomedical Informatics, Columbia University Irving Medical Center, New York, NY 10032; JP Sulzberger Columbia Genome Center, Columbia University, New York, NY 10032; Program for Mathematical Genomics, Columbia University Irving Medical Center, NY 10032

**Keywords:** protein fitness landscape, protein language model, self-supervised learning, sequence likelihood, mutation effect

## Abstract

Protein language models, and models that incorporate structure or homologous sequences, estimate sequence likelihood *p(sequence)* that reflects the protein fitness landscape and is commonly used in mutation effect prediction and protein design. It is widely believed in deep learning field that larger model performs better across tasks. However, for fitness prediction, language model performance declines beyond a certain size, raising concerns about their scalability. Here, we showed that model size, training dataset, and stochastic elements can bias the predicted *p(sequence)* away from real fitness. Model performance on fitness prediction depends on how well *p(sequence)* matches evolutionary patterns in homologs, which is best achieved at a moderate *p(sequence)* level for most proteins. At extreme predicted wild-type sequence likelihoods, models predict uniformly low or high likelihoods for nearly all mutations, failing to reflect the real fitness landscape. Notably, larger models tend to predict proteins with higher *p(sequence)*, which may exceed the moderate range and thus reduce performance. Our findings clarify the scaling behavior of protein models on fitness prediction and provide practical guidelines for their application and future development.

## Introduction

Characterizing the protein fitness landscape—how mutations affect protein function, abundance, activity, and interaction—is a central challenge in biology. It is crucial for elucidating the mechanisms underlying diseases, advancing precision medicine, guiding viral surveillance, and advancing protein design and engineering. While deep mutational scanning^1^ (DMS) experiment has been applied to measure mutation effects on diverse proteins^2,3^, it’s time-consuming, labor-intensive, and limited to molecular effects that are easy to assay. To complement experimental efforts, supervised machine learning methods have been developed by training on curated mutation datasets^4–6^. However, these datasets are limited in size and biased toward functionally important genes^4^, restricting the models’ generalizability and robustness^7^. As a result, there is an urgent need for predictive models capable of estimating the protein fitness landscape without training on curated mutation datasets.

In recent years, self-supervised models have been developed for zero-shot fitness prediction by estimating sequence likelihood *p(sequence)* (Throughout this manuscript, “sequence likelihood” and *p(sequence)* refer to the likelihood of wild-type sequence unless otherwise noted for mutant sequences.): the probability of a protein sequence under the learned distribution of natural proteins. Self-supervised models often match or even surpass the performance of supervised models^8,9^. Representative models include protein language models (pLMs) such as ESM2^10^; multi-sequence alignment (MSA)-based models like MSA-Transformer^11^ and EVE^8^; inverse folding models such as ESM-IF1^12^, which predict sequence from structure; and hybrid models like ESM3^13^, which integrate sequence, structure, and other information. Based on their training data, these methods fall into two main categories: general models trained on massive datasets of tens to hundreds of millions of proteins from diverse protein families, and family-specific models trained on MSA of individual protein family (e.g., EVE, which requires training a separate model for each MSA). MSA-Transformer integrates both family-specific information and general information across diverse protein families by being trained on millions of MSAs^11^.

These models are trained to maximize the likelihood of training protein sequences using strategies such as masked or next token prediction^14^, conditioned on sequence, structure, or MSA (**Figure 1A**). While some models incorporate additional training objectives^13^, only sequence prediction is used to predict fitness (see Methods for details)^3^. Fitness of a mutation is estimated by comparing the predicted likelihoods of the mutant and wild-type sequences^3,9^ (**Figure 1A**), usually by calculating the log-likelihood ratio (LLR). In this framework, an LLR close to zero means that the mutation is nearly as fit as the wild-type, implying a neutral mutation effect, whereas a strongly negative LLR indicates the mutation is much less fit and potentially deleterious. For models trained via masked prediction, computing the *p(sequence)* is intractable. Instead, pseudo-likelihood and LLR calculated from marginal approaches are used (see Methods for details)^3,9^.

**Figure 1.**
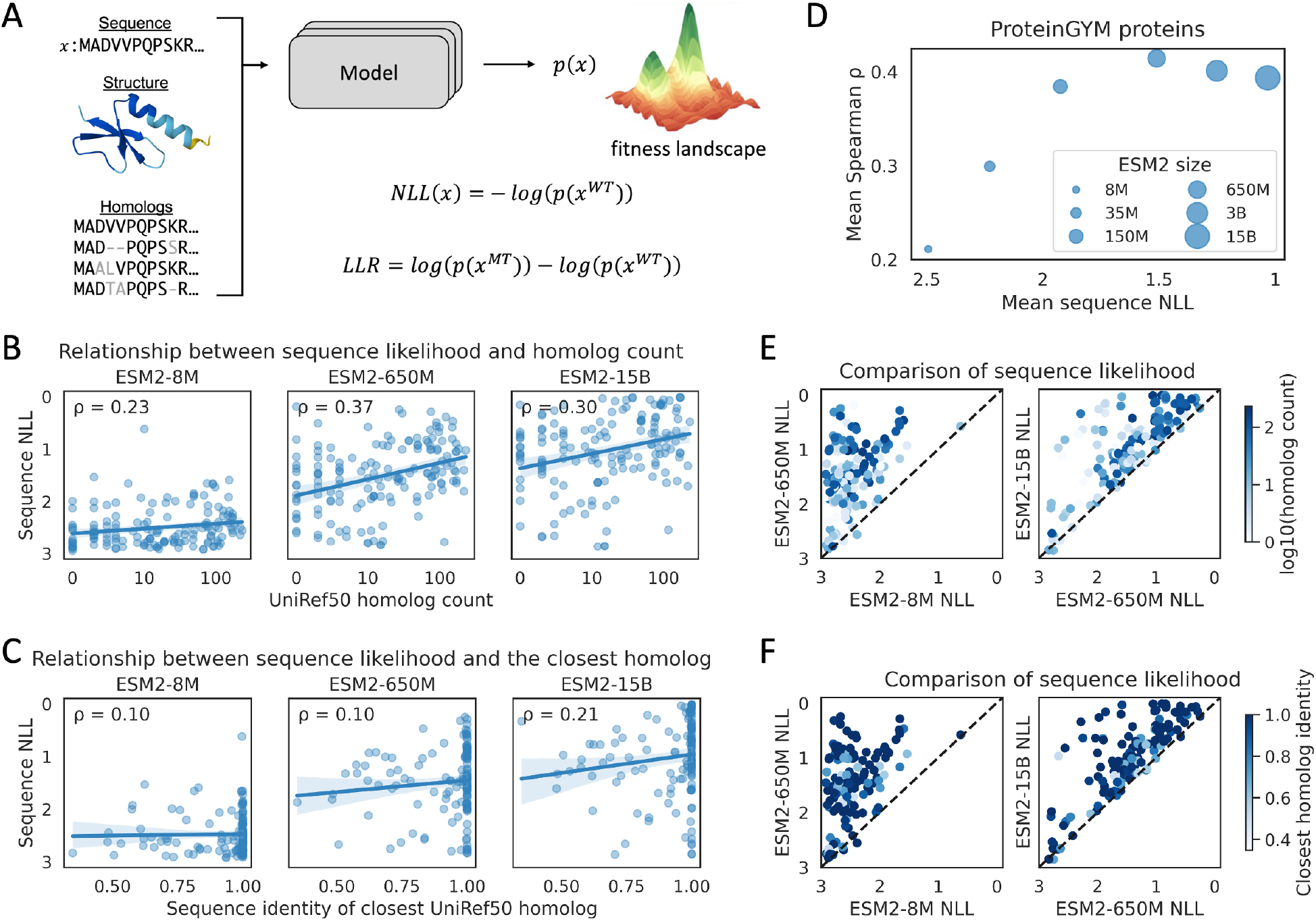
Model-predict sequence likelihood is influenced by factors unrelated to protein fitness. **A**, Overview of the calculation of sequence likelihood and log-likelihood ratio (LLR). Models are trained to predict sequence likelihood *p*(*x*) using information from sequence, structure, and homologs, negative log-likelihood (NLL) is usually used as the training loss. The lower-left panel shows a multiple sequence alignment (MSA), the dashed lines indicate alignment gaps, and the grey-shaded amino acids denote positions in homologous sequences that differ from the query sequence (the first sequence in the MSA). For mask-prediction-based models, pseudo-likelihood is used. LLR is calculated based on the entire sequence for generative models, and masked residues for mask-prediction-based models. **B**, Relationship between predicted sequence likelihood and the number of homologs. Each point represents a protein; the x-axes show the number of UniRef50 homologs in the log scale. Homologs are defined as those with ⩾20% sequence identity and ⩾80% coverage. The curves represent linear regressions, with the shaded areas indicating the 95% confidence intervals. ρ represents Spearman correlation. **C**, Relationship between predicted sequence likelihood and sequence identity of the closest homolog. **D**, The weighted mean performance on fitness prediction, and mean predicted sequence likelihood of ESM2 models on proteins in 154 ProteinGYM experiments. Point size indicates ESM2 model size. The performance is first averaged within each of the five functional categories in ProteinGYM, and the weighted mean performance is computed as the average of five category-specific performances. These values are reported in Table S1. **E-F**, Comparison of sequence likelihoods predicted by different ESM2 models. Each point represents a protein; colors indicate the log-scaled number of homologs (**E**) and sequence identity of the closest homolog (**F**).

Among these models, pLMs gain particular attention for their strong performance and minimal input requirement—they only require sequence as input, without needing MSA or structure. In deep learning field, scaling up models is a common strategy to improve performance on downstream tasks^15^. pLM scaling has proven effective for masked or next residue prediction and structure modeling^10,16^; however, for fitness prediction, model performance declines beyond a certain size^3,17^: ESM2-650M (million parameters) and 3B (billion parameters) outperform the 15B model (**Table S1**), xTrimoPGLM-3B^18^ outperforms the 10B and 100B models, and autoregressive pLMs ProGen3-1B and 3B outperform the 10B and 46B models^16^. This raises a key question in model development: why does scaling up pLMs not consistently improve performance on fitness prediction? While pLMs are typically released in multiple sizes, enabling direct investigation of the scaling behavior, other models are usually released in one size, making it unclear whether the scaling trend applies to them. Beyond scaling, several important questions remain: Under what conditions are these models most effective? How to choose appropriate models for a given protein?

In this study, we systematically investigated the relationship between sequence likelihood and the performance of fitness prediction across diverse models. We found that various factors unrelated to fitness—including model size, training dataset, and stochastic elements—can influence general model-predicted likelihoods, making them uninformative for fitness prediction in extreme cases. By analyzing large fitness benchmarks^2,3^, we found that the performance of general models depends on how well their predicted *p(sequence)* aligns with evolutionary patterns in homologs. Notably, they perform best at a moderate *p(sequence)* level for most proteins, which explain the scaling behavior of pLMs as medium-sized models like ESM2-650M predict more proteins with moderate *p(sequence)*. Our findings clarify how likelihood-based self-supervised models predict fitness and lay the groundwork for developing next-generation fitness predictors.

## Results

### General model-predicted *p(sequence)* is influenced by factors unrelated to protein fitness

As general models trained on large datasets function as black boxes, we examined whether factors unrelated to fitness can influence the predicted *p(sequence)*. We first investigated two key factors in scaling up models: the size of the training dataset and the number of trainable parameters. To do this, we analyzed predictions from several models on 154 fitness measurements from deep mutational scanning (DMS) experiments in the ProteinGYM^3^ benchmark (see Methods for data filtering), with a primary focus on six ESM2 models spanning 8 million to 15 billion parameters. ProteinGYM includes proteins from diverse taxa and five functional categories, spanning mutation effects on activity, binding, stability, organismal fitness, and expression.

Larger training datasets include more protein families and more homologs within each family. Since dissimilar proteins have little influence on the predicted likelihood of a protein^19^, we focused on the role of its homologs. We identified UniRef50^20^ (the ESM2 training dataset, version2021_04) homologs with over 20% sequence identity and 80% coverage to proteins in the ProteinGYM benchmark. By analyzing the number of homologs, we found that proteins with more homologs tend to have higher predicted likelihoods (**Figure 1B**, quantified using negative log-likelihood (NLL, *–log p(sequence)*)). We note that this trend is not indirectly driven by conservation level differences of protein families (**Figure S1A**). However, proteins with high predicted likelihoods do not necessarily have many UniRef50 homologs (**Figure 1B**). We then analyzed the sequence identity of the closest homologs and observed that proteins with high predicted likelihoods often have highly similar homologs in UniRef50. But the presence of such homologs does not guarantee high predicted likelihoods (**Figure 1C**). We also analyzed the summed sequence identity × coverage of all homologs, which provides a more quantitative measure of the overall similarity of homologs in the training set. The results are similar to those obtained using homolog count (**Figure S1B**). Additionally, we observed similar relationships between homologs in the training set and sequence likelihoods for SaProt^21^, a structure-informed model (**Figure S2A-B**). Overall, homologs in the training set can influence model-predicted likelihoods, but the relationship is complex.

We then analyzed model size. These models are typically trained using the NLL of masked or next token as the loss function. As larger models achieve lower training loss, they tend to predict higher sequence likelihoods^10,16,18^. We observed this trend for the proteins in the ProteinGYM benchmark (**Figure 1D**). Furthermore, we examined the magnitude of likelihood increase as model size scaled up and observed substantial variability: some proteins showed little or no increase, while others exhibited large gains (**Figure 1E-F**). Notably, the magnitude of likelihood increase between larger and smaller models is not clearly associated with the likelihoods from the smaller model, nor with the number or similarity of homologs in the training set (**Figure 1E-F**). We observed similar results for SaProt, xTrimoPGLM^18^, and ProGen3^16^ (**Figure S2C-D**).

These results suggest that, beyond the homologs in training dataset and model size, additional factors influence model-predicted likelihoods. One such factor could be the stochastic elements introduced during model training, including parameter initialization, data shuffling, and masked residue sampling. To directly assess the impact of these stochastic elements, we analyzed five ESM1v^22^ models, which share the same architecture and training dataset but were trained with different random seeds. We found that approximately 10% of ProteinGYM proteins show NLL differences greater than 0.5 among five ESM1v models (**Figure S3**), and the maximum observed difference is 1.2: the protein Mafg (UniProt ID: O54790) has a predicted sequence NLL of 1.1 in ESM1v_5 but 2.3 in ESM1v_2. This result suggests that stochastic elements can lead models to converge on different local minima, resulting in variability in predicted sequence likelihoods for some proteins. Overall, model size, training dataset, stochastic elements, and other unknown factors complicate the interpretation of general model-predicted likelihoods.

### Magnitude of predicted *p(sequence)* affects fitness estimation by influencing LLR values

As model-predicted sequence likelihood is affected by many factors, the LLR value, directly tied to sequence likelihood (LLR is the NLL difference between wild-type and mutant sequences, **Figure 1A**) and used to estimate fitness, is also affected. This may impact fitness prediction performance. To investigate this, we focused on model size, as other factors are difficult to control given models we have access to. Larger models, achieving lower training loss, tend to assign higher probability to the wild-type amino acid and lower probabilities to the others (since the total probability per site sum to one), resulting in LLRs with larger magnitudes. In extreme cases, a non-informative model that predicts equal probability to all 20 amino acids yields LLRs of zero for all mutations, while an overconfident model that assigns a probability of one to the wild-type and zero to all others produces LLRs of negative infinity. Although certain proteins may predominantly harbor neutral or deleterious mutations, the collapsed LLR distributions in these two extreme scenarios are uninformative for most proteins.

To illustrate this, we examined predictions of six ESM2 models on PTEN (phosphatase and tensin homolog), one of the most extensively studied proteins, for which DMS experiments of both cell growth^23^ and protein stability^24^ are available. Across both DMS experiments, the distribution of mutation effects is clearly bimodal, with approximately 20% of mutations exhibiting deleterious effects (**Figure 2A**). However, the distributions of ESM2 predicted LLR vary substantially with model size. Smaller models predict LLRs clustered near zero, while larger models predict strongly negative LLRs to most mutations (**Figure 2B, S4A**), mirroring the two extreme scenarios we described above. Notably, the medium-sized model ESM2-150M reproduces the bimodal distribution observed in experiments (**Figure 2A, S5A**) and achieves the best performance (**Figure S5B**), with Spearman correlations of 0.55 for growth and 0.46 for stability. In contrast, both the smallest (ESM2-8M) and largest (ESM2-15B) models yield correlations below 0.3 in both DMS experiments (**Figure 2B, S4A, S5B**). We also observed that xTrimoPGLM-1B model best captures the distribution of experimental mutation effects and outperforms larger models with 3B to 100B parameters (**Figure S5C-D**). We note that the smallest available xTrimoPGLM model is 1B, and thus comparisons to smaller models are not possible. We further investigated this relationship at the residue level by comparing the mean mutation effect per residue with the predicted probability of the wild-type residue. Residues assigned high probability by the model tend to have strongly negative LLRs for mutations, indicating that such residues are mutation sensitive. ESM2-15B assigns high probability approaching one to nearly all residues, while ESM2-8M assigns high probability to very few residues— both failing to capture the experimentally observed distribution of mutation-sensitive residues (**Figure 2C-D, S4B**). In contrast, ESM2-150M, predicts residue probability that better reflect mean mutation effects (**Figure 2C-D, S4B**). Additionally, we analyzed PTEN cancer hotspot mutations^25^, and found that predicted residue likelihood from medium-sized ESM2 models (35M and 150M) outperform other ESM2 models at identifying residues with hotspot mutations (**Figure S5E**). These results suggest that medium-sized ESM2 models more accurately capture the residue-level importance in PTEN. We observed similar results in other proteins that also exhibit a rise-then-fall performance trend with increasing ESM2 model size (**Figure S4C-H**).

**Figure 2.**
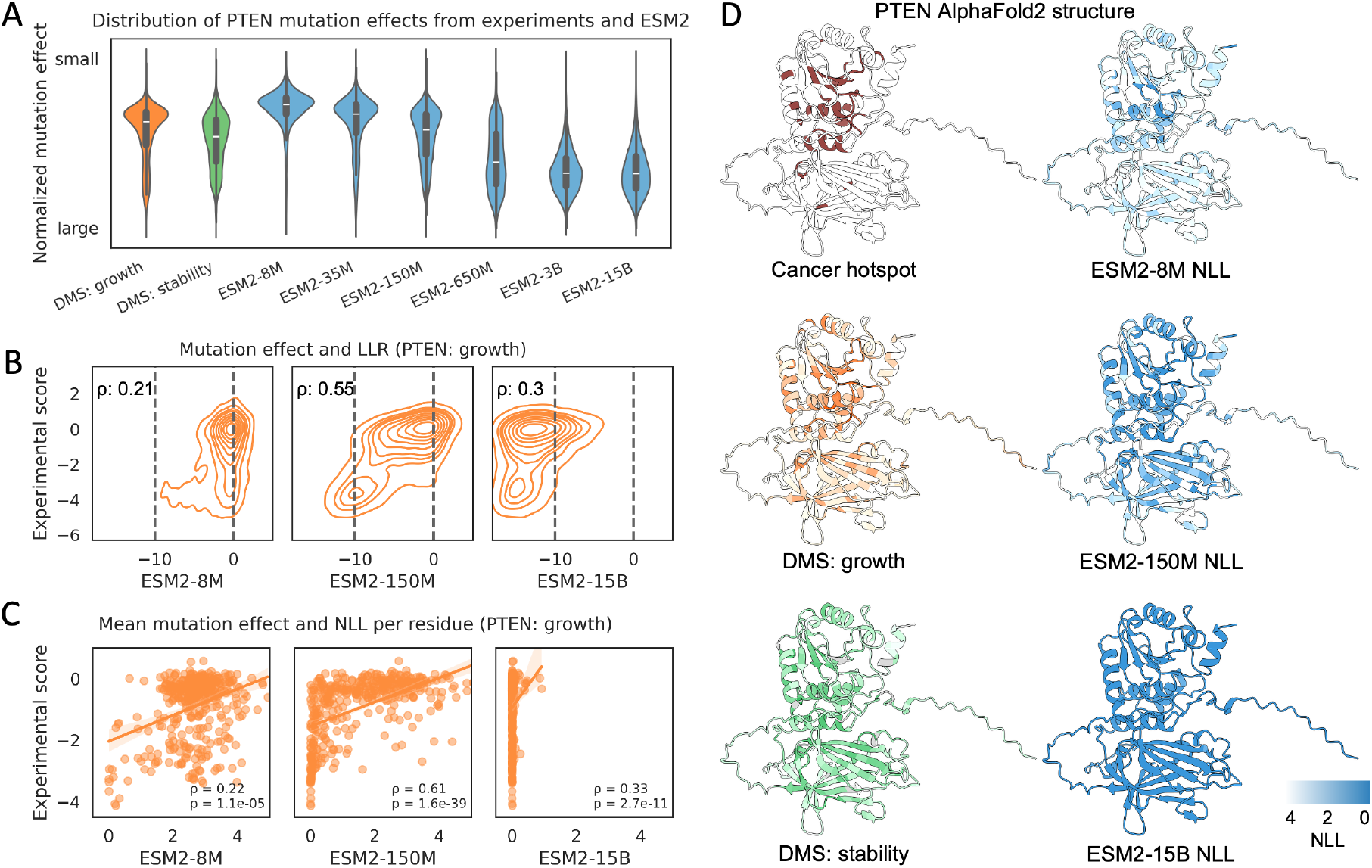
Distributions of experimental and predicted PTEN mutation fitness. **A**, Distributions of normalized mutation effects from DMS experiments and ESM2 predictions. Experimental effects and ESM2 LLRs are normalized to the range of 0–1 for visualization. ESM2 LLRs are calculated using the masked marginal approach. **B**, Relationship between ESM2-predicted LLRs and experimental effects for PTEN mutations. The x-axis represents predicted LLR, the y-axis represents log-scaled, wild-type–normalized fitness measurements based on the growth of humanized yeast. ρ: Spearman correlation. **C**, Relationship between ESM2-predicted probability per residue (quantified using NLL) and mean experimental effects for mutations at each residue (residues with at least 10 mutations are shown). The x-axis represents predicted NLL per residue, the y-axis represents mean experimental scores per residue from ProteinGYM, each point represents a residue. Spearman correlation and corresponding p-values are shown. **D**, Visualization of mutation sensitivity and ESM2-predicted NLL per residue on the AlphaFold2-predicted PTEN structure. For two DMS experiments, the mean mutation effect at each residue is shown (grey color indicates no mutation at the site). Darker colors indicate stronger mutation effects or higher ESM2 predicted likelihood (i.e., lower NLL).

### General model performance on fitness prediction peaks at a moderate level of *p(sequence)*

The above analyses highlight a critical caveat when using general models for fitness prediction: their predicted likelihoods can be influenced by unrelated factors and may not reliably reflect real fitness. To systematically investigate the relationship between model-predicted sequence likelihood and performance on fitness prediction, we evaluated models on 154 DMS experiments from the ProteinGYM benchmark^3^ (see Methods for dataset filtering).

We first evaluated six ESM2 models. Although larger models consistently predict higher sequence likelihood (**Figure 1D**), higher sequence likelihood does not always lead to better performance. Notably, the ESM2-650M model outperforms the larger 3B and 15B models (**Figure 1D, Table S1**). By comparing wild-type sequence likelihoods with performances of 154 experiments, we observed a bell-shaped relationship using the non-parametric LOWESS^26^ modeling (locally weighted scatterplot smoothing, black curves in **Figure 3**), with performance peaking at a moderate likelihood level—corresponding to a wild-type sequence NLL of approximately 1.2 (**Figure 3**, *p(sequence)* ≈ 0.3). Because second-order polynomial regressions closely match the non-parametric LOWESS trends (**Figure 3**), we also used it to describe the bell-shaped relationship and reported its 95% confidence intervals. Notably, within the optimal likelihood range, all ESM2 models, regardless of size, perform comparably (**Figure 3**). We observed the same trend for ESM2 models on the mega-scale protein folding stability dataset^2^ (**Figure S6A**), indicating that this relationship is not specific to ProteinGYM. These results indicate that the level of sequence likelihood, rather than model size, is the primary determinant of model performance on fitness prediction. While more homologs in the ESM2 training set can lead to higher predicted sequence likelihoods (**Figure 1B**), we found that the performance drop at high likelihood is not caused by homolog overrepresentation in the training set (**Figure S1C**, see **Figure S2F** for SaProt).

**Figure 3.**
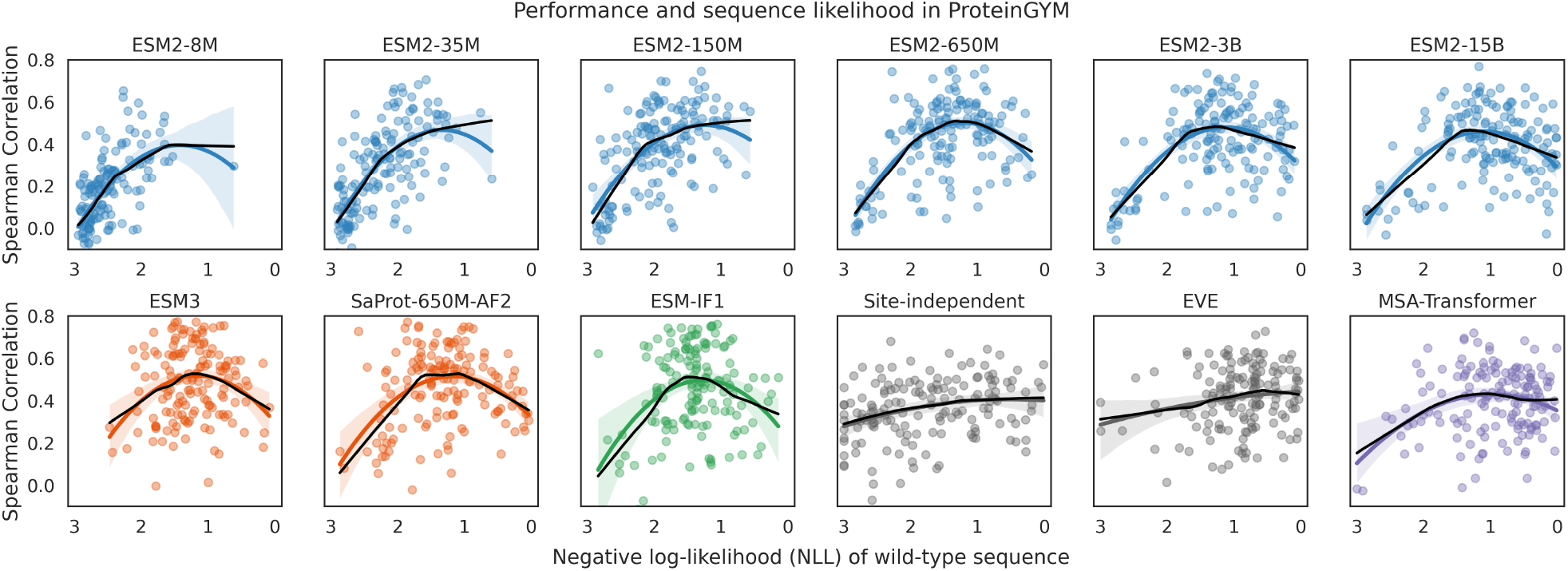
Relationship between fitness prediction performance and model-predicted wild-type sequence likelihood. The y-axes show the Spearman correlation between model predicted LLRs and experimental effects, the x-axes represent the negative log-likelihood (NLL) of wild-type sequences. LLRs of ESM2, ESM3, SaProt, and MSA-Transformer are calculated using the masked marginal approach. LLRs of ESM-IF1 and EVE are calculated using the full sequence (see Methods for details). The LLR of the site-independent model is computed as the log difference in amino acid frequency at each site. Only residues with mutations are included in the NLL calculation. Each point represents an experiment from ProteinGYM (154 experiments after filtering). The black curves show LOWESS modeling, while the colored curves show second-order polynomial regressions, with shaded areas indicating the 95% confidence intervals, all regression analyses were performed using the Python package Seaborn. Colors indicate model types: blue: protein language models; orange: hybrid models, green: inverse-folding models, grey: family-specific models trained on MSA of individual protein family, and purple: MSA-Transformer that integrate both general and family-specific information.

We further evaluated a broad range of general models that predict fitness using sequence likelihood. These included Transformer encoder–based mask language models ESM1v^22^ and ESMC^27^, whose training datasets are different from that of ESM2; the convolution-based model CARP^28^; and Transformer decoder– based generative models ProGen3^16^ and RITA^29^. We also evaluated structure-sequence-hybrid models ESM3^13^, ProSST^30^, and SaProt^21^, as well as the inverse folding model ESM-IF1^12^ (see Methods for details). Despite differences in architecture, input modalities, and training strategies, all these general models exhibit the bell-shaped relationship between performance and wild-type sequence likelihood (**Figure 3, S7A**; see Discussion for decoder-based pLMs). Remarkably, peak performances are achieved at the similar level of sequence likelihood (**Figure 3, S7A**). For ProSST and small ESM2 models, only one side of the bell-shaped trend is observed, as almost no proteins exhibit predicted sequence likelihoods beyond the optimal range due to limited model capacity. Because ProSST shares an almost identical model design with SaProt, we expect it to exhibit the same bell-shaped relationship. In addition, for all LLR-based models, fitness predictive power necessarily vanishes when the wild-type sequence NLL is zero: in this limit, models predict LLRs of negative infinity to all mutations. Consequently, the performance curve must pass through the origin (NLL = 0, fitness prediction Spearman correlation = 0). In practice, however, sequence NLL values of zero are not observed for these models. We used the masked marginal approach for models trained via masked token prediction. We note that we also observed the bell-shaped trend when these models are applied using the wild-type marginal approach (**Figure S7B-D**, see Methods for details).

Beyond general models trained on diverse protein families, we also analyzed family-specific models. These included simple frequency-based site-independent models that treat each position independently, with or without homologous sequence weighting (see Methods for details), and the variational autoencoder-based model EVE^8^, which captures inter-residue dependencies. These models are trained independently on MSA of each protein family with no or less parameters, making them less susceptible to the unrelated factors that affect general models. Notably, these family-specific models do not exhibit the bell-shaped relationship between performance and sequence likelihood (**Figure 3, S7A**). For MSA-Transformer^11^ which integrates both general and family-specific information, the bell-shaped trend is present but less pronounced (**Figure 3**).

### The bell-shaped relationship arises from models’ varying ability to capture context information

Fitness prediction requires models to capture both context information (i.e., the sequence and structural context that determine mutation sensitivity) and substitution specificity (i.e., how well different amino acids fit a given context). To investigate the origin of the bell-shaped relationship between performance and sequence likelihood, we disentangled two components. Context understanding is quantified by comparing the mean LLR and the mean experimental effect for mutations on each residue within a protein. Substitution specificity is assessed by correlating LLRs with experimental effects of 19 mutations at each residue.

By comparing model performance on the two components with predicted likelihood, we found that general models exhibit the bell-shaped relationship between context understanding (mean mutation effect prediction performance) and wild-type sequence likelihood, with performance peaking at a similar likelihood range (**Figure 4A, S8A, S6B**). Extreme cases are exemplified by our results for PTEN and other proteins (**Figure 2C, S4**). In contrast, per-residue performance increases monotonically with predicted likelihood (i.e., the predicted probability of each residue; **Figure 4B, S8B, S6C**; see Discussion for explanation). Family-specific models, however, do not display the bell-shaped trend for either component (**Figure 4, S8**). Notably, all models show substantially stronger performance in understanding context than substitution specificity on 154 experiments (**Figure 4, S8**). None of the models evaluated in ProteinGYM achieves mean Spearman correlation above 0.3 for substitution specificity. These findings indicate that the fitness predictive power of current models primarily stems from their ability to capture protein context, which also underlies the bell-shaped relationship observed between fitness prediction performance and sequence likelihood in general models.

**Figure 4.**
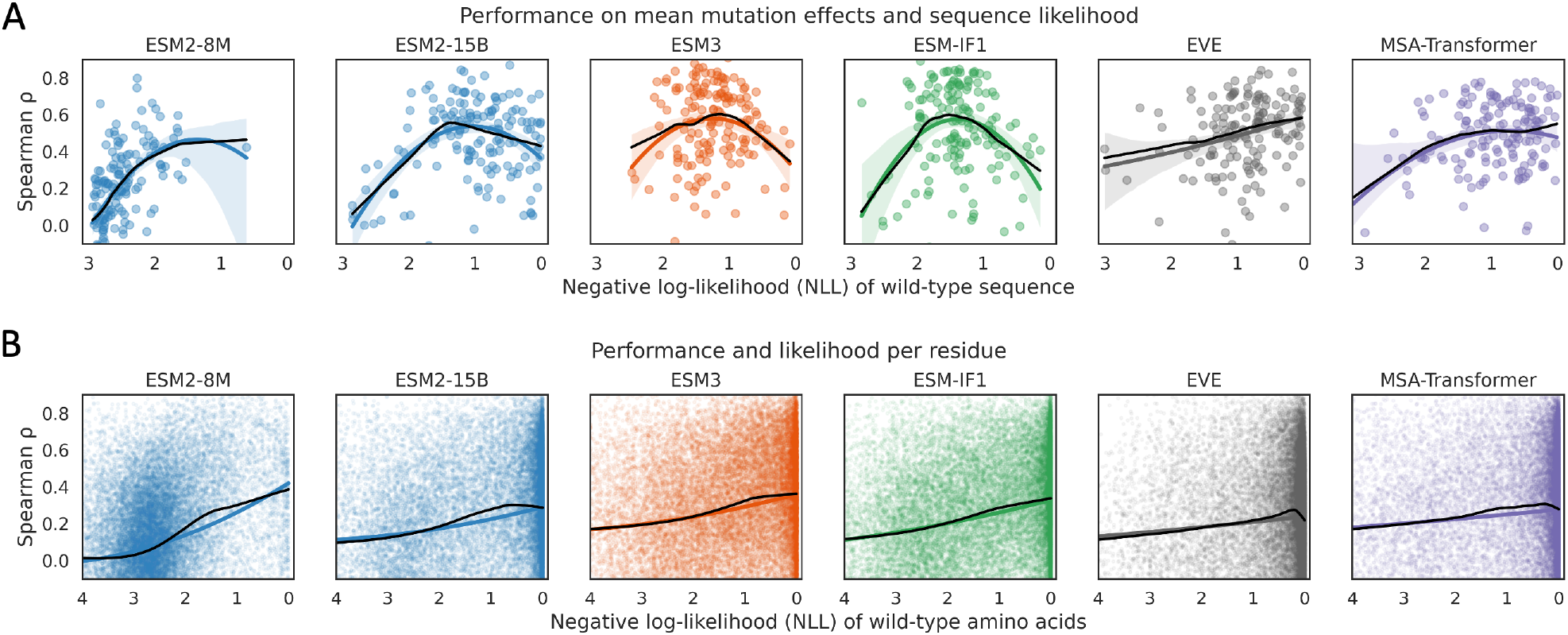
Model performance on mean mutation effects and mutation effects per residue. **A**, The y-axes show the Spearman correlation between mean LLRs and mean experimental mutation effects per residue in each protein, reflecting model understanding of protein context. The x-axes represent the NLL of wild-type sequences. Only residues with mutations are included in the NLL calculation. Each point represents a ProteinGYM experiment. **B**, The y-axes show the Spearman correlation between LLRs and experimental effects of all 19 mutations per residue, reflecting model understanding of substitution specificity. The x-axes represent the negative log predicted probability of each residue. Each point represents a residue. The black curves show LOWESS modeling, while the colored curves show second-order polynomial regressions, with shaded areas representing the 95% confidence intervals.

### Comparing model-predicted likelihood to biophysical context and mutation sensitivity

To describe the context of each residue more quantitatively, we considered both the biophysical aspect and mutation sensitivity. Biophysical context is measured using relative solvent accessibility (RSA), which is closely associated with mutation effects^31,32^: mutations at buried sites tend to destabilize the protein and lead to loss of function, whereas mutations at surface residues have relatively small effects on stability on average, with a fraction involved in functions. We observed correlations between RSA and the mean mutation effect per residue in ProteinGYM experiments, with stronger correlations observed for experiments of stability (**Figure 5A**). Although the mean experimental mutation effect reflects mutation sensitivity of each residue, it is not comparable across DMS experiments. Therefore, we manually examined the distribution of experimental effects. Among the 154 experiments, 122 display bimodal distributions (as exampled in **Figure S9A**). For these experiments, we manually defined thresholds to distinguish deleterious and neutral mutations. The thresholds were determined by visual inspection to identify the separation point between two modes in the distribution of mutation effects, and the corresponding threshold values are provided in the Supplementary Data 1. Mutation sensitivity at each site was then quantified by counting the number of deleterious mutations among the 19 possible mutations. We observed a strong inverse relationship between RSA and the number of deleterious mutations (**Figure 5B; S9B**), supporting the connection between protein structure and mutation sensitivity (functional importance).

**Figure 5.**
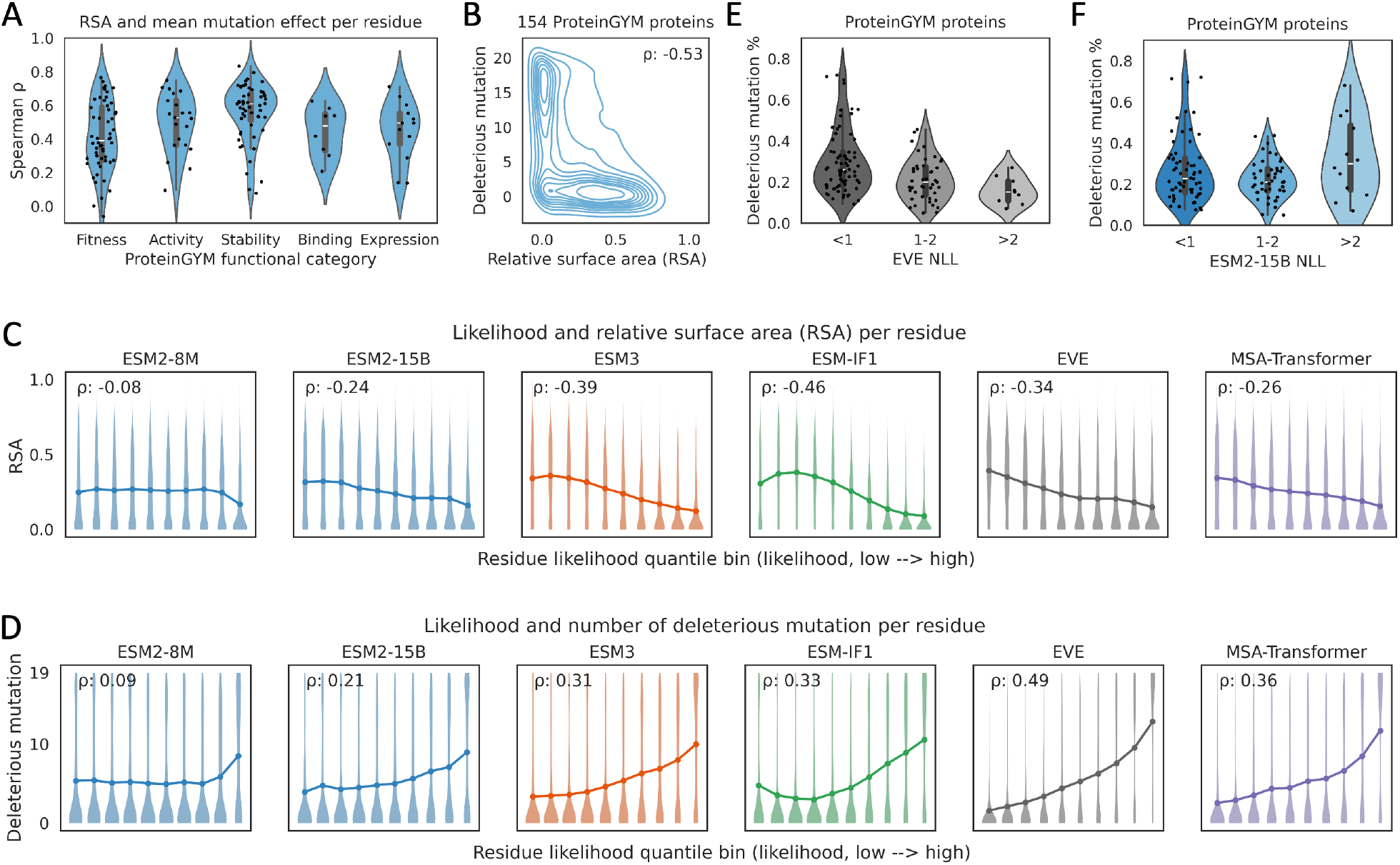
Comparison of model-predicted residue likelihood with the biophysical context and mutation sensitivity. **A**, Correlations between relative solvent accessibility (RSA) and mean mutation effect per residue. Each point represents a ProteinGYM experiment. **B**, The relationship between RSA and number of deleterious mutations per residue. 122 experiments showing bimodal distributions of mutation effect are analyzed. **C-D**, Residues are grouped into ten quantile bins based on model-predicted probability. For each bin, the distribution and mean value are shown for RSA (**C**, range 0–1) and the number of deleterious mutations per residue (**D**, range 0–19). Only results for experiments with bimodal distribution of mutation effect are shown, only residues with 19 mutations are shown. **E–F**, 122 DMS experiments were classified into three groups based on sequence NLL from EVE or ESM2-15B. The proportion of deleterious mutations in each protein is shown.

By comparing model-predicted per-residue likelihoods with the biophysical context and mutation sensitivity, we found that residues with high predicted likelihoods by all methods tend to have lower RSA and more deleterious mutations (**Figure 5C-D**). This demonstrates that both general and family-specific models can capture biophysical and functional context to some extent. Biophysical context is better captured by structure-informed models such as ESM3 and ESM-IF1 at high likelihoods (**Figure 5C**), owing to their direct use of structural input. Mutation sensitivity, however, is better captured by family-specific models (**Figure 5D, S10**). Among the top 10% of high-likelihood residues, the mean number of deleterious mutations is 8.5 for ESM2-8M, 9.0 for ESM2-15B, 9.9 for ESM3, and 10.5 for ESM-IF1, compared to 12.8 for EVE and 11.7 for MSA-Transformer. Conversely, among the bottom 10% of low-likelihood residues, the mean number of deleterious mutations is 5.3 for ESM2-8M, 3.9 for ESM2-15B, 3.3 for ESM3, and 4.9 for ESM-IF1, compared to 1.6 for EVE and 2.5 for MSA-Transformer (**Figure 5D**). Furthermore, at the protein level, we observed that proteins with high EVE sequence likelihood tend to harbor more deleterious mutations (**Figure 5E**)—a trend not observed for ESM2 (**Figure 5F**). These results indicate that general model predicted likelihood fails to reflect functionally importance (mutation sensitivity) as reliably as that of family-specific models.

### The performance of general models depends on how well predicted *p(sequence)* matches evolutionary patterns in homologs

Finally, we set out to explain the bell-shaped relationship between sequence likelihood and fitness prediction performance. We began by comparing the predicted likelihoods from different methods. Family-specific model predicted likelihoods directly reflect evolutionary patterns in homologous sequences: a high residue-level likelihood indicates a conservated residue, while a high protein-level likelihood indicates a large fraction of conserved residues. Even though the family-specific models we evaluated employ different architectures, their predicted likelihoods are correlated at both the residue and protein levels (**Figure 6A-B**). General models, in contrast, implicitly learn evolutionary information from large-scale datasets. Likelihoods predicted by different general models are also correlated at both the residue and protein levels (**Figure 6A-B**). When comparing general models with family-specific models, correlations are observed at the residue level (**Figure 6A**), suggesting that general models capture the correct overall trend. However, for protein level likelihoods, general models show weak or no correlation with family-specific models (**Figure 6B**; decoder-based pLMs show weak protein-level correlations, see Discussion).

**Figure 6.**
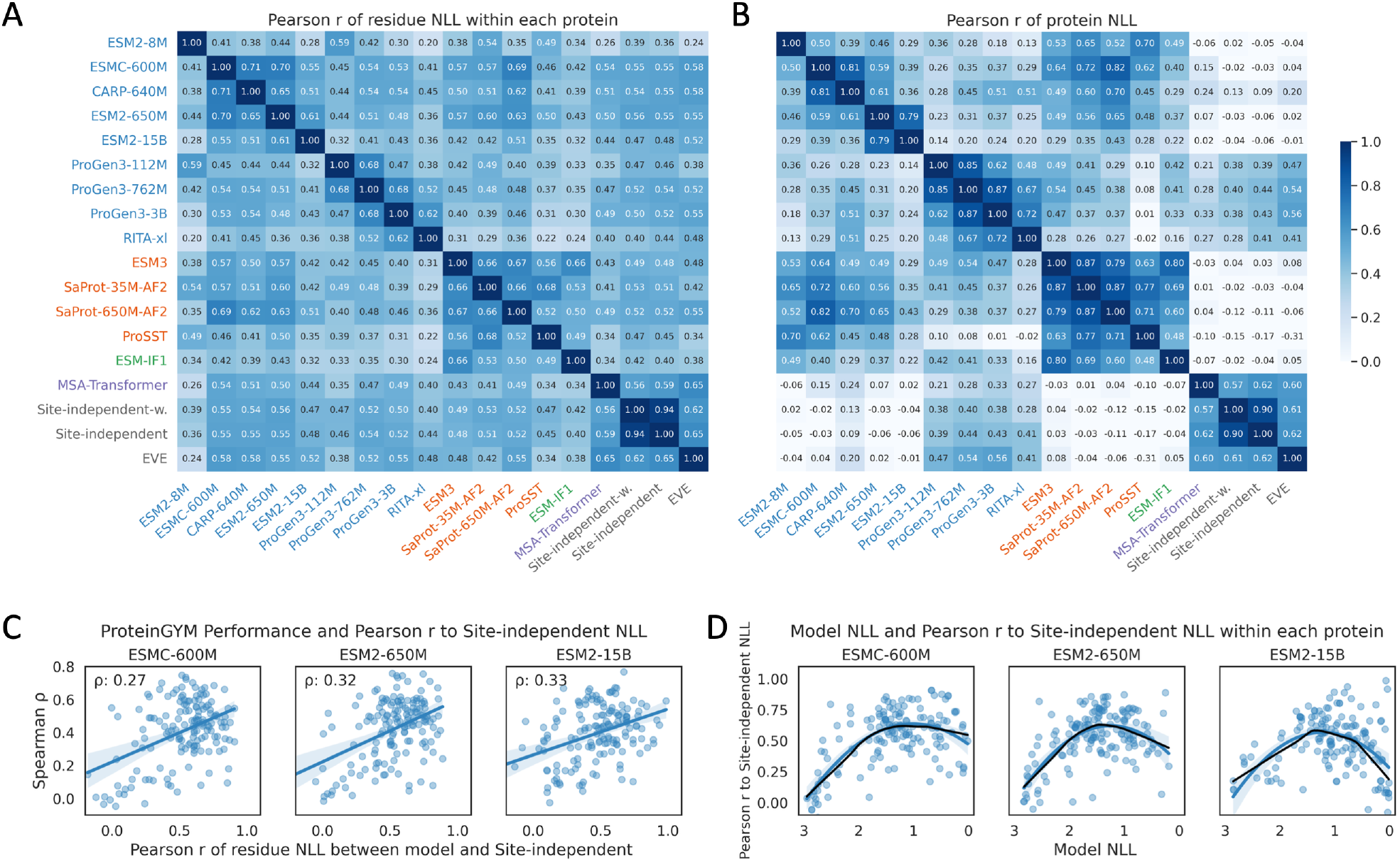
The performance of general models depends on how well they capture evolutionary information. **A**, Pearson correlation of per-residue NLL within each protein, the mean correlations across proteins in 154 experiments are shown. **B**, Pearson correlation of whole-sequence NLL, which is calculated as the mean of per-residue NLL within the protein. **C**, Relationship between ESM fitness prediction performance and per-residue NLL correlation to the site-independent model. The y-axes show fitness prediction performance. The x-axes show the Pearson correlation between ESM NLL and site-independent NLL per residue within each protein. Each point represents an experiment. The curves represent linear regressions, with the shaded areas indicating the 95% confidence intervals. **D**, Relationship between ESM sequence likelihood and correlation to site-independent NLL. The y-axes show the Pearson correlation between ESM NLL and site-independent NLL per residue within each protein. The x-axes show the ESM2 NLL. Each point represents an experiment. The black curves show LOWESS modeling, while the colored curves represent second-order polynomial regressions, with the shaded areas indicating the 95% confidence intervals.

This may be because the per-residue NLL correlation is only moderate (Pearson r ≈ 0.5; **Figure 6A**), and aggregating residue-level likelihoods to the protein level amplifies noise. This explains why ESM2-predicted likelihoods capture mutation sensitivity at the residue level (**Figure 5D**) but fail to reflect the proportion of deleterious mutations at the protein level (**Figure 5F**). Because likelihoods derived from homologs by family-specific models better reflect protein fitness (**Figure 5D-E**), we therefore hypothesized that the bell-shaped curve arises because general models capture evolutionary information most effectively when their predicted sequence likelihood falls within a moderate range.

To test this, we compared ESM models to family-specific models: the site-independent model, EVE, and MSA-Transformer. These models capture evolutionary patterns from sequence alone. We used likelihoods from family-specific models to represent evolutionary patterns in homologous sequences and quantified how well ESM models capture evolutionary patterns by computing the Pearson correlation between their predicted residue-level likelihoods. We found that the better ESM models capture evolutionary information, the better they perform in fitness prediction (**Figure 6C, S11A**). Notably, ESM models capture evolutionary information most strongly at moderate sequence likelihoods (**Figure 6D, S11B**), which corresponds to the range where their fitness prediction performance is optimal (**Figure 3**). These results are consistent across the three family-specific models we evaluated (**Figure 6C-D, S11A-B**). Therefore, the ability of ESM models to predict fitness relies on how well predicted likelihoods align with evolutionary patterns. These findings suggest that we should first assess whether the predicted sequence likelihoods align with evolutionary patterns, derived from family-specific models, before applying general models to fitness prediction. Considering both the fraction of proteins exhibiting this alignment and the resulting performance gains relative to no filtering, we recommend using the per-residue NLL Pearson correlation around 0.5 as the cutoff (ProteinGYM in **Figure S11C**, protein stability dataset^2^ in **Figure S11D**).

We also compared other general models to family-specific models and found that ESM3 and ESM-IF1 behaved differently (**Figure S12**). Unlike sequence-only pLMs, these structure-informed models can capture not only evolutionary patterns derived from homologs but also biophysical constraints (**Figure 5C**), the latter being particularly useful for modeling mutation effects on protein stability (**Table S1, Figure S13**).

## Discussion

Predicting the protein fitness landscape is one of the most important applications of pLMs and other deep learning models in biology. Our work explains why these models can predict fitness, why larger models do not always perform better, and under what conditions they should or should not be used for fitness prediction. Overall, we found that the better a general model captures evolutionary information, the better its performance on fitness prediction. However, unlike family-specific models that directly utilize evolutionary patterns in homologs, general models are influenced by unrelated factors which can decouple predicted likelihoods from true evolutionary patterns. This disconnect leads to poor performance on fitness prediction at extreme sequence likelihood, which leads to the bell-shaped relationship between model performance and predicted likelihood. The optimal likelihood for general models corresponds to *p(sequence)* ≈ 0.3. Interestingly, a 30% sequence identity threshold is commonly used to search homologous sequences^33^. One reason for the bell-shaped relationship is the probabilistic coupling between wild-type and mutant residues imposed by the SoftMax operation in these models. Because supervised models usually do not use sequence likelihood to infer mutation effects and do not enforce this probabilistic coupling, they are not expected to exhibit this scaling behavior. Supervised fine-tuning of pLMs for fitness prediction showed that larger models tend to achieve better performance^16^.

Besides the factors we mentioned in the result section, model-predicted sequence likelihood is also influenced by model architecture and training strategy. For example, models that use MSAs as input tend to predict higher likelihoods, as it’s easier to predict sequence with homologous sequences as input. Training strategies also play a role: higher weights on the regularization loss can suppress overfitting, reducing likelihoods of training sequences. Moreover, sequence patterns can also affect predicted likelihoods^34^. Therefore, predicted likelihoods should be interpreted with caution. In our evaluation, most analyses were conducted within each model, so differences in model architecture and training strategy do not affect our results. For comparisons between models, we used Pearson correlation which captures similarity in likelihood patterns independent of absolute values.

We observed a monotonic relationship between model predicted residue likelihood and performance on 19 mutations at each residue (**Figure 4B**), we explained it as follows: residues assigned low probabilities correspond to positions where general models poorly capture contextual information or where family-specific models suffer from poor homolog alignment. This results in noisy predicted probabilities of the 20 amino acids and reduces per-residue fitness prediction performance. In contrast, residues with higher predicted probabilities indicate that general models better capture contextual information, or that family-specific models have good homolog alignment. As shown in Figure 5D, these residues tend to correspond to functionally important positions. Figure 6A further suggests that general models identify these residues by capturing the conservation signal. For these residues, models assign high probability to the wild-type amino acid and low probability to alternatives, causing most mutations to be predicted as deleterious. This aligns with their functional importance (mutation sensitive) and results in improved per-residue performance.

The bell-shaped relationship is less pronounced for the decoder-based autoregressive pLMs ProGen3 and RITA compared with other general models (**Figure S7A**), and their peak performance at the optimal likelihood remains relatively low. This likely reflects an intrinsic limitation of autoregressive pLMs for fitness prediction: they cannot simultaneously incorporate contextual information from both upstream and downstream residues (ESM-IF1 is also autoregressive, but it leverages complete structural context). Nonetheless, autoregressive models have advantages over masked models: they can explicitly estimate sequence likelihood, whereas masked language models rely on pseudo-likelihood; and they are trained on all residues of training sequences, while masked models are trained only on a subset of residues masked during training. These may explain why autoregressive pLMs show correlations with family-specific models in protein level likelihood, whereas masked models do not (**Figure 6B**). Additionally, autoregressive models are suited for generating protein sequences of varying lengths.

Weinstein et al.^35^ studied the benefits of model misspecification in fitness prediction and attributed the decreased performance of larger pLMs to their improved density estimation. Here, we show that likelihoods predicted by pLMs do not always reliably reflect the true density of homologs (**Figure 6B**); thus, the reduced performance of larger pLMs cannot be solely explained by improved density estimation. Gordon et al.^36^ reported that the preference (measured by likelihood) for a given protein sequence established during pretraining is predictive of pLM fitness prediction performance. However, they did not provide a mechanistic explanation. Yu et al.^37^ found that model prediction entropy is related to viral protein fitness prediction, with low entropy (corresponding to high sequence likelihood) associated with better performance. Gurev et al.^38^ reported that larger pLMs perform better on viral protein fitness prediction. While their conclusions on viral proteins may appear to conflict with our findings, we note that viral proteins are underrepresented in protein datasets and are therefore assigned relatively low likelihoods by general models (**Figure S14A**). As a result, their sequence likelihoods have not reached the optimal levels observed for well-represented proteins, thus the expected performance decline with increasing likelihood or model size is not observed (**Figure S14A**). This phenomenon also extends to other proteins that are poorly learned by general models, such as de novo designed proteins (**Figure S14B**).

Understanding model-predicted likelihood is a fundamental question in language models and the key to understand how models generate their outputs. Interestingly, a bell-shaped relationship has also been observed between LLM sentence likelihoods and human quality judgments^39^. Thus, an important caveat for both LLMs and protein models is that data points predicted with high likelihoods may not be real or biologically meaningful. Prior work has shown that overparameterized models trained on small datasets tend to memorize data: assigning high likelihoods to training data. In contrast, smaller models trained on large datasets tend to generalize by learning shared patterns^40^. For applying protein models to mutation effect prediction, certain level of generalization, where the model can integrate information from homologs, is more desirable. In our analysis, current models appear to operate in the intermediate (determined by the ratio of training tokens to model parameters^40^): they memorize some proteins / residues, generalize to a subset, and ignore others. Which proteins / residues a model chooses to memorize, generalize, and ignore remains an open question.

Our results offer practical guidance for applying general models to predict fitness, we recommend first verifying whether the predicted likelihoods align with evolutionary patterns in homologs, which can be calculated using simple family-specific models (**Figure S11C-D**). For training next generation fitness predictors, we recommend incorporating evolutionary patterns during training. This can be achieved by estimating evolutionary patterns from homologs beforehand and encouraging alignment between predicted likelihoods and evolutionary patterns within the training objective. While our study focuses on protein mutation effects, the issue we found is general and may extend to DNA/RNA language models and other applications, such as protein design.

## Supplementary information

Supplementary Figures S1-14

Supplementary Table S1

Supplementary Date:

1. 154 DMS experiments evaluated in this study, including manually defined cutoffs for classifying deleterious and neutral mutations.
2. Performance on 154 DMS experiments across different methods.
3. Mean mutation effect prediction performance.
4. Performance per residue.
5. Predicted likelihood per residue from different methods.

## Author contributions

Y.S. and C.H. conceived the study. C.H. performed the analyses and drafted the manuscript. Y.S. and C.H. interpreted the results. All authors revised the manuscript.

## Competing interests

The authors declare no competing interests.

## Acknowledgment

This work was supported by NIH grants R35GM149527 and Simons Foundation SFARI #1019623.

## Data and code availability

All mutation data used in this manuscript are publicly available from ProteinGYM (https://proteingym.org/) and Zenodo (DOI: 10.5281/zenodo.7401274). The codes for running models are the same as that provided by ProteinGYM (https://github.com/OATML-Markslab/ProteinGym). The site-independent models and marginal approaches can be readily implemented following the method descriptions in our work.

## Methods

### Dataset, protein structure, and sequence analysis

In this study, we focused exclusively on single-residue substitutions. Other mutation types—including multiple-residue substitutions, insertions, deletions, and truncations—were not considered for the following reasons: (1) current models’ performance on these mutations is markedly lower compared to single substitutions^3^; (2) several of the evaluated methods cannot be directly applied to these mutation types; and (3) while some studies have extended models to these mutation types, the predictions are derived from prediction of single-residue substitutions^9^ (e.g., the prediction of a multiple-residue substitution is calculated as the sum of the corresponding single-substitution predictions, and the prediction of a truncation is calculated as the maximum single-substitution prediction in the truncated region).

Experimental mutation effects, model predictions, Multiple sequence alignments (MSAs), and protein structures were downloaded from the ProteinGYM^3^ website (https://proteingym.org/) and its GitHub repository (https://github.com/OATML-Markslab/ProteinGym) in May 2025. From the 217 mutational scanning datasets available, only residues with experimental effects of all 19 possible substitutions were included. Restricting to proteins with at least 20 such residues resulted in a dataset comprising 486,932 single-residue substitution mutations across 154 experiments (Supplementary Data 1). We note that this ProteinGYM filtering criterion was applied to all analyses except for the PTEN analysis, where we included residues with at least ten measured substitutions to increase residue coverage. The thresholds for classifying deleterious and neutral mutations provided in ProteinGYM are not used, as they are defined based on the distribution of mutation effects of both single and multiple substitutions and appear unreasonable for some proteins in our visualizations. Instead, we manually defined thresholds using only single substitutions. The cutoffs were determined by visual inspection to identify the separation point between the two modes in the distribution of experimental single-mutation effects. Cutoffs were defined only for DMS experiments exhibiting a clear bimodal separation, as exampled in Figure S9A. The corresponding cutoff values are provided in Supplementary Data 1 to ensure reproducibility.

The mega-scale protein folding stability dataset^2^, including both designed and natural mini domains, was analyzed in Figure S4E-H, S6, S11D, and S14B. Single-residue substitutions in the table “Tsuboyama2023_Dataset2_Dataset3_20230416.csv” were analyzed. For applying the site-independent model to these proteins in Figure S11D, homologs were identified using the ColabFold^41^ server with default parameters, only homologs with less than 50% gaps were analyzed.

PTEN cancer hotspot mutations were downloaded from cBioPortal^25^ (February 2026). The curated set of non-redundant studies was used, missense mutations annotated as “CancerHotspot” or “3DHotspot” were included.

ProteinGYM provides predictions from a wide range of methods, only likelihood-based methods were evaluated here. For models (EVE, MSA-Transformer, ESM1v) offering both single-model and ensemble predictions, the single-model predictions were used. Protein solvent-accessible surface areas (SASA) were calculated using the MDTraj^42^ package (version: 1.10.2). Relative surface areas were computed by dividing the SASA of each residue by the maximum SASA observed for that amino acid across all proteins. For homolog detection, MMseqs2^43^ (version 16.747c6) was used to search against UniRef50^20^ (version 2021_04, which was used to train ESM2) with the following parameters: *--min-seq-id 0*.*2 -c 0*.*8 --max-accept 1000*.

### Applying masked language models for protein fitness prediction

Masked language models (MLMs) evaluated in the study include ESM2^10^, ESM1v^22^, ESMC^27^, CARP^28^, ESM3-open^13^ (1.4B parameters), SaProt^21^, ProSST^30^ (110M parameters, 2048 structure tokens), and MSA-Transformer^11^. Among them, ESM2, ESM1v, ESMC, and CARP are sequence-only MLMs; ESM3, ProSST, and SaProt are sequence and structure hybrid MLMs; MSA-Transformer is an MSA-based MLM. For MLM, computing the exact sequence likelihood *p*(***x***) is intractable. Instead, pseudo-likelihood is used, where each residue is masked individually and predicted in turn. For a protein of length *L* with sequence ***x*** *=* (*x*_1_, …, *x*_*L*_) (*x*_*i*_ denotes the wild-type amino acid, which is also explicitly written as *x*_*i, wt*_ when it needs to be distinguished from mutations in the following sections.), structure ***s*** (structure token of the protein is also a list of tokens with length *L*), and a set of homologs ***h***, the pseudo-likelihood of sequence-only MLM is defined as:

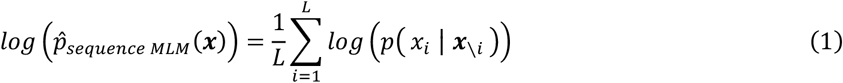

Where ***x***_*∖i*_ denotes the sequence with position *i* masked.

ESM3, ProSST, and SaProt were trained with both sequence and structure tokens. ESM3 predicts the probabilities of sequence and structure tokens separately, ProSST only predicts masked sequence during training, whereas SaProt predicts them jointly. Specifically, SaProt predicts the probabilities of 400 combinations of sequence–structure tokens, corresponding to 20 amino acids × 20 FoldSeek^44^ structure tokens. During fitness inference, all structure tokens were provided, while the residue at the mutation site was masked. For SaProt, the predicted probability for each amino acid was obtained by summing over 20 sequence–structure combinations with that amino acid. The pseudo-likelihood of sequence and structure hybrid MLM is defined as:

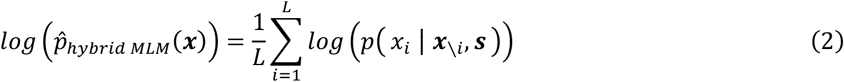

MSA-Transformer leverages homologous sequences to provide evolutionary context. The choice of homologs can be made by random sampling, selecting the most similar, or selecting the most dissimilar sequences^11^. In this study, we followed the protocol used in ProteinGYM, where homologs were sampled according to sequence-similarity–based weights (also used for EVE). The pseudo-likelihood of MSA-Transformer is defined as:

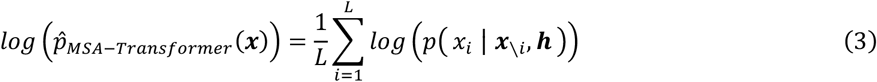

For fitness prediction, the log-likelihood ratio (LLR) between the mutated sequence ***x***_***mt***_ and the wild-type sequence ***x***_***wt***_ can be computed using the pseudo-likelihood of their full sequences:

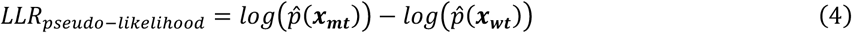

However, this approach is computationally intensive, since for each mutated sequence of length *L*, computing the pseudo-likelihood requires *L* forward passes through the model, making it impractical for large-scale fitness inference. Instead, simplified approximations are used, including the masked marginal and wild-type marginal approaches, both of which consider only the predicted probabilities at the mutation site rather than the likelihood of the entire sequence. The underlying hypothesis is that the difference in predicted probabilities at a single residue is proportional to the difference in overall sequence likelihood.

For the masked marginal approach, the residue at the mutation site is masked, and the model directly predicts the probabilities of the wild-type and mutant amino acids, the LLR is then calculated as:

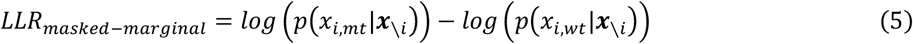

Previous studies have shown that for MLMs trained with the standard masking scheme (e.g., ESM2, where 15% of tokens are randomly selected during pre-training, with 80% replaced by the mask token, 10% kept unchanged, and 10% replaced by a random token), predictions from the unmasked wild-type sequence can be used directly for fitness inference^9^. In this case, the LLR is calculated as:

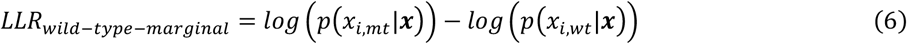

The computational cost differs substantially among these methods. For a protein of length *L*, scoring all possible single-residue substitutions (19 per position) requires running the model *L*×*19*×*L* times with ^*LLR*_*pseudo*−*likelihood*_, *L* times with *LLR*_*masked* − *marginal*_, and only one time with *LLR*_*wild* − *type* − *marginal*_. In this^ study, we used masked marginal approach for all MLM methods, which is the most used methods to estimate fitness from MLM. For MLMs trained with a fraction of tokens substituted and a fraction of tokens unchanged, the sequence likelihood computed without masking is highly correlated with that from the masked approach^19^ (Figure S7B), and the LLR scores derived from both approaches show nearly identical performance in predicting mutation effects^9^ (Figure S7C). Therefore, our conclusions also apply when using MLMs with the wild-type marginal approach (Figure S7D).

For all models, the predicted probability of all possible tokens in each position sum to one:

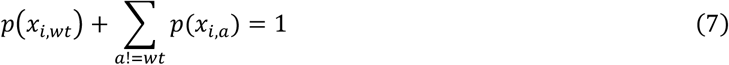

Where *a* represent amino acids different from the wild-type. ProteinGYM provides LLRs for the MLMs we evaluated, but not the underlying sequence likelihoods. However, given the definition of LLRs (Eq. 5) together with Eq. 7, the predicted residue probabilities can be inferred using the following equations:

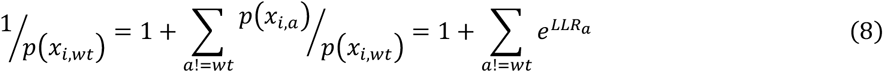

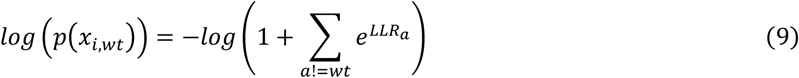

Eq. 8 and 9 were used to estimate sequence likelihoods for positions with LLRs of all 19 possible mutations in the ProteinGYM dataset, eliminating the need to run these models. As ProteinGYM and the default ProSST implementation use the wild-type marginal approach, masked marginal scores were computed for ProSST to ensure comparability of sequence likelihoods across methods. Sequence likelihoods of five ESM1v models and ESM2 prediction for the mega-scale protein folding stability dataset^2^ were computed by us.

### Applying autoregressive models for protein fitness prediction

We evaluated several autoregressive generative models: pLMs RITA-xl^29^ (1.2B parameters) and ProGen3^16^, and the inverse-folding model ESM-IF1^12^. Both RITA and ProGen3 are decoder-only transformer models, with ProGen3 incorporating mixture-of-experts layers. These models were trained in an autoregressive manner: RITA and ProGen3 on both forward and reverse sequences, and ESM-IF1 on forward sequences only (N-terminal to C-terminal). For autoregressive models, the likelihood of the forward sequence can be computed explicitly as:

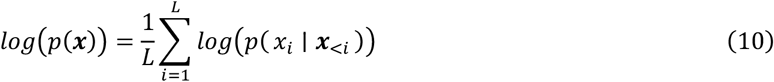

Where ***x***_<*i*_ denotes the residues before position *i*. For autoregressive pLMs, the likelihood of a sequence was computed as the mean of predictions from the forward and reverse sequences:

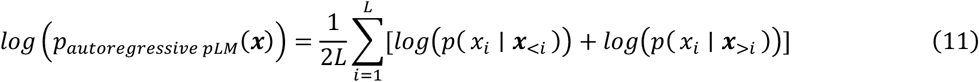

For ESM-IF1, the likelihood of a sequence was computed as:

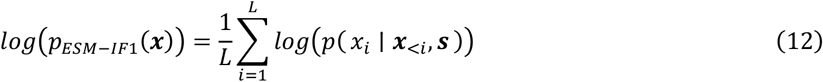

For fitness prediction using autoregressive models, the sequence likelihood is computed directly for each mutated sequence. Since the wild-type sequence likelihood is constant, calculating LLR is unnecessary for performance evaluation. ProteinGYM provides negative log likelihood (NLL) for each mutated sequence rather than LLRs. The sequence likelihoods for wild-type proteins were computed by us using Eq. 11 and 12.

We also investigated whether autoregressive pLMs could be applied using marginal approaches, which can reduce the times of model running from 2× the number of mutant sequences to only two (forward and reverse wild-type sequences). In this setting, the probabilities of both the wild-type and mutant amino acids at a given position are predicted using upstream and/or downstream residues as context. Using ProGen3-3B, we evaluated four strategies: (1) computing the LLR from the forward sequence only, (2) computing the LLR from the reverse sequence only, (3) averaging the LLRs obtained from the forward and reverse sequences, and (4) averaging the predicted probabilities from the forward and reverse sequences before computing the LLR. The results are summarized in Table S1. Performances of all marginal approaches are substantially lower than using the ProGen3-3B full sequence likelihood (**Table S1**). Thus, in the Results section, only the results obtained using the full sequence likelihood for generative pLMs were reported. We note that the marginal approaches for autoregressive pLMs also exhibit weak bell-shaped relationships between wild-type sequence likelihood and fitness prediction performance (data not shown).

### Site-independent models

The site-independent model was used as the baseline family-specific model. MSAs provided by ProteinGYM were used, and homologous sequences with more than 50% gaps were removed. For each site, the probabilities of the 20 amino acids and the gap character were calculated as:

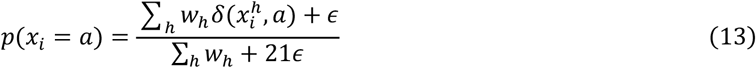

where *w*_*h*_ denotes the weight of homologous sequence *h*, which was either calculated using the same strategy as in EVE or set as one for all sequences. *δ* is the Kronecker delta. *ϵ* is the pseudo count, for the unweighted model, *ϵ* was set to 1; for the weighted model, *ϵ* was set to the minimum positive value in the frequency table. Fitness was estimated using LLR of mutant and wild-type amino acids at each position. Benchmarking on the ProteinGYM mutations evaluated in this study, both the site-independent (unweighted) and the site-independent-weight model achieve mean Spearman correlations around 0.36 (**Table S1**). Despite their simplicity, they outperform the site-independent model trained with EVmutation^45^ (mean Spearman correlation 0.33).

### Family-specific model EVE and its marginal approach

EVE^8^ models were trained separately for each MSA using the code provided in the ProteinGYM GitHub repository, taking approximately a week on ten A6000 GPUs. Model was trained with the random seed of 0 and *threshold_focus_cols_frac_gaps* = 1 (all positions were modeled). As a variational autoencoder^46^, EVE does not provide exact sequence likelihoods. Instead, the evidence lower bound (ELBO) was used as a proxy, calculated as:

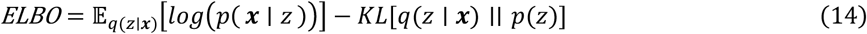

Where *z* is the latent variable, *q*(*z* ~ ***x***) is the approximate posterior from the encoder, *p*(***x*** ~ *z*) is the decoder likelihood, and *KL*[~~] denotes the Kullback–Leibler divergence (KLD). The ELBO was computed for both wild-type and mutated sequences by averaging over 20,000 samples of *z*. The decoder conditional probability *p*(***x*** ~ *z*) was used to approximate the probability of each amino acid at each site, averaged over the 20,000 samples using wild-type sequence as input.

Using EVE to estimate fitness is computationally intensive, as it requires training a separate model with millions of parameters for each MSA, and during inference, the decoder must be run tens of thousands of times per mutated sequence to obtain a stable ELBO estimation. To reduce this burden, we implemented a simplified approach, which we termed EVE-marginal. In this approach, the conditional probability *p*(***x*** ~ *z*) is obtained from the wild-type sequence only, and fitness is estimated like the MLM wt-marginal approach. EVE-marginal performs comparable to the full EVE model (**Table S1**), while being substantially efficient.

## Notes

### Competing Interest Statement

The authors have declared no competing interest.

### Summary of Updates

updated main figures 1-6 and supplementary figures during the revision in a journal. also added more analyses to support our conclusion

