## Supplemental table and figures for "Understanding Language Model Scaling on Protein Fitness Prediction"

Hou et al.

Table S1
Figure S1-14

**Table S1: Performance (Spearman correlation) summary on ProteinGYM selected assays.**

| Model class | Model name | Organismal Fitness (56) | Stability (56) | Activity (21) | Expression (13) | Binding (8) | Average of five functions | Average of 154 assays |
|---|---|---|---|---|---|---|---|---|
| pLM (masked) | ESM2-8M | 0.117 | 0.191 | 0.258 | 0.226 | 0.263 | 0.211 | 0.18 |
| | ESM2-35M | 0.192 | 0.326 | 0.376 | 0.291 | 0.311 | 0.299 | 0.28 |
| | ESM2-150M | 0.304 | 0.427 | 0.45 | 0.37 | **0.37** | 0.384 | 0.378 |
| | ESM2-650M | 0.383 | **0.445** | **0.489** | **0.393** | 0.362 | **0.414** | **0.42** |
| | ESM2-3B | 0.382 | 0.424 | 0.477 | 0.39 | 0.331 | 0.401 | 0.408 |
| | ESM2-15B | 0.38 | 0.39 | 0.46 | 0.391 | 0.346 | 0.393 | 0.394 |
| | ESM1v | 0.367 | 0.332 | 0.461 | 0.375 | 0.327 | 0.373 | 0.366 |
| | ESMC-600M | 0.366 | 0.432 | 0.477 | 0.385 | 0.359 | 0.404 | 0.407 |
| | CARP-640M | **0.39** | 0.368 | 0.445 | 0.36 | 0.298 | 0.372 | 0.382 |
| pLM (autoregressive) | RITA-xl (1.2B) | 0.379 | 0.267 | 0.405 | 0.374 | 0.338 | 0.353 | 0.34 |
| | ProGen3-112M | 0.274 | 0.188 | 0.36 | 0.26 | 0.304 | 0.277 | 0.255 |
| | ProGen3-762M | 0.366 | 0.27 | **0.43** | 0.347 | 0.345 | 0.352 | 0.337 |
| | ProGen3-3B | **0.393** | **0.315** | 0.41 | **0.403** | **0.347** | **0.374** | **0.365** |
| | ProGen3-3B-marginal (forward) | 0.307 | 0.117 | 0.329 | 0.289 | 0.183 | 0.245 | 0.233 |
| | ProGen3-3B-marginal (reverse) | 0.336 | 0.173 | 0.318 | 0.317 | 0.249 | 0.279 | 0.268 |
| | ProGen3-3B-marginal (forward+reverse 1) | 0.35 | 0.223 | 0.373 | 0.333 | 0.238 | 0.303 | 0.3 |
| | ProGen3-3B-marginal (forward+reverse 2) | 0.336 | 0.168 | 0.363 | 0.313 | 0.233 | 0.283 | 0.271 |
| Structure-informed models | ESM3 (open, 1.4B) | 0.376 | **0.613** | 0.446 | 0.456 | 0.443 | 0.467 | 0.482 |
| | SaProt-35M-AF2 | 0.284 | 0.53 | 0.432 | 0.421 | 0.399 | 0.413 | 0.411 |
| | SaProt-650M-AF2 | 0.369 | 0.529 | **0.495** | 0.473 | 0.405 | 0.454 | 0.455 |
| | ESM-IF1 | 0.329 | 0.595 | 0.406 | 0.425 | 0.415 | 0.434 | 0.449 |
| | ProSST (110M, K=2048) mask | **0.393** | 0.589 | 0.465 | **0.490** | **0.482** | **0.484** | **0.487** |
| MSA-based models | Site-independent | 0.38 | 0.326 | 0.397 | 0.371 | 0.343 | 0.363 | 0.36 |
| | Site-independent-weight | 0.373 | 0.315 | 0.392 | 0.366 | 0.345 | 0.358 | 0.353 |
| | EVE | **0.44** | 0.365 | 0.475 | **0.422** | **0.383** | **0.417** | **0.413** |
| | EVE-marginal | 0.425 | **0.374** | **0.48** | 0.41 | 0.348 | 0.407 | 0.408 |
| | MSA-Transformer | 0.42 | 0.352 | 0.467 | 0.407 | 0.356 | 0.4 | 0.397 |

Marginal approaches were explored for generative models, ProGen3 and EVE, to reduce computational cost (see Methods for details). For ProGen3-3B, four marginal strategies were evaluated: log-likelihood ratios (LLRs) calculated from the forward sequence only, LLRs calculated from the reverse sequence only, averaged LLRs from the forward and reverse sequences (forward+reverse 1), and averaging predicted probabilities from the forward and reverse sequences followed by LLR calculation (forward+reverse 2). The marginal strategy for EVE is described in the Methods.

Although a monotonic trend in performance was observed for the evaluated ProGen3 models, we note that larger ProGen3 models with 10B and 46B parameters were reported to underperform the 3B model in the original ProGen3 study; these larger models are not publicly available and are therefore not evaluated here.

The site-independent and site-independent-weight models are described in the Methods.

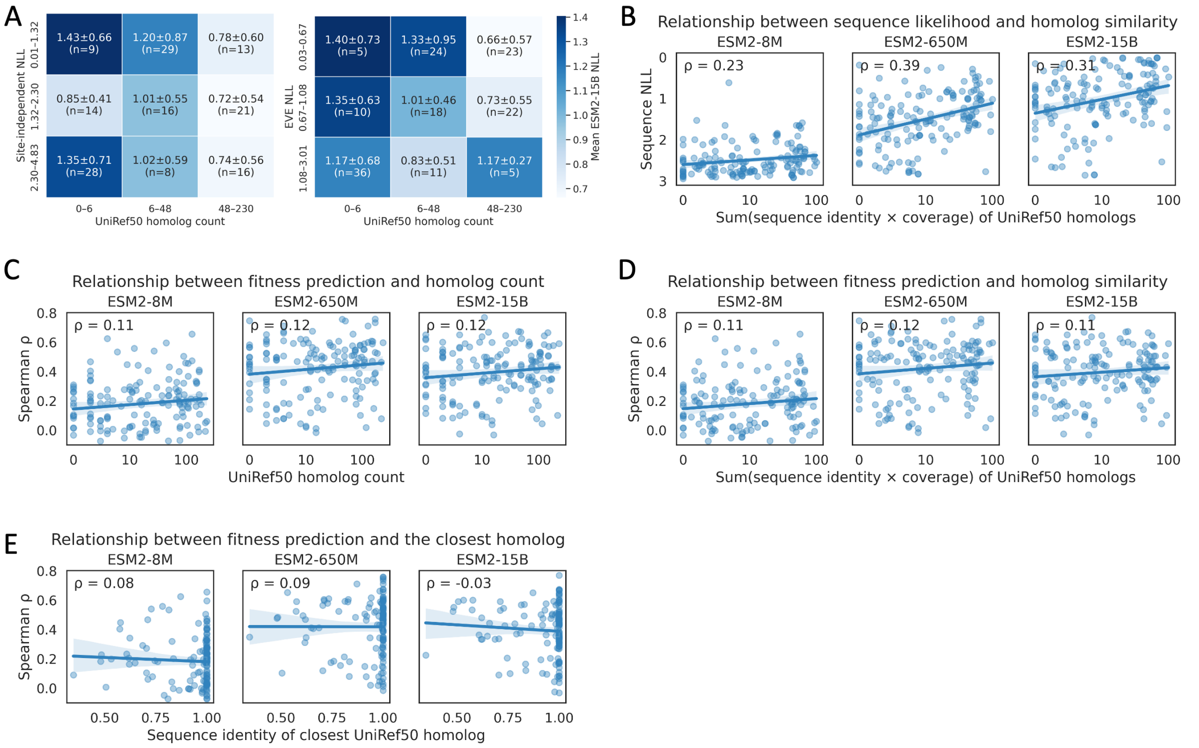


**Figure S1. Impact of homologs in training set on ESM2 sequence likelihood and fitness prediction performance.** **A**, Proteins in 154 ProteinGYM assays are stratified into nine groups defined by three conservation levels (sequence likelihood from family-specific models) and three homolog-count levels. The mean and standard deviation of sequence NLL from ESM2-15B are shown. **B**, Relationship between predicted sequence likelihood and the similarity of all homologs. The similarity is computed as the sum of (sequence identity × alignment coverage) across all homologs in the training data. Each point represents a protein; the x-axes show values in the log scale. **C-E**, Relationship between fitness prediction performance and homologs in UniRef50. The curves represent linear regressions, with the shaded areas indicating the 95% confidence intervals. ρ represents Spearman correlation.

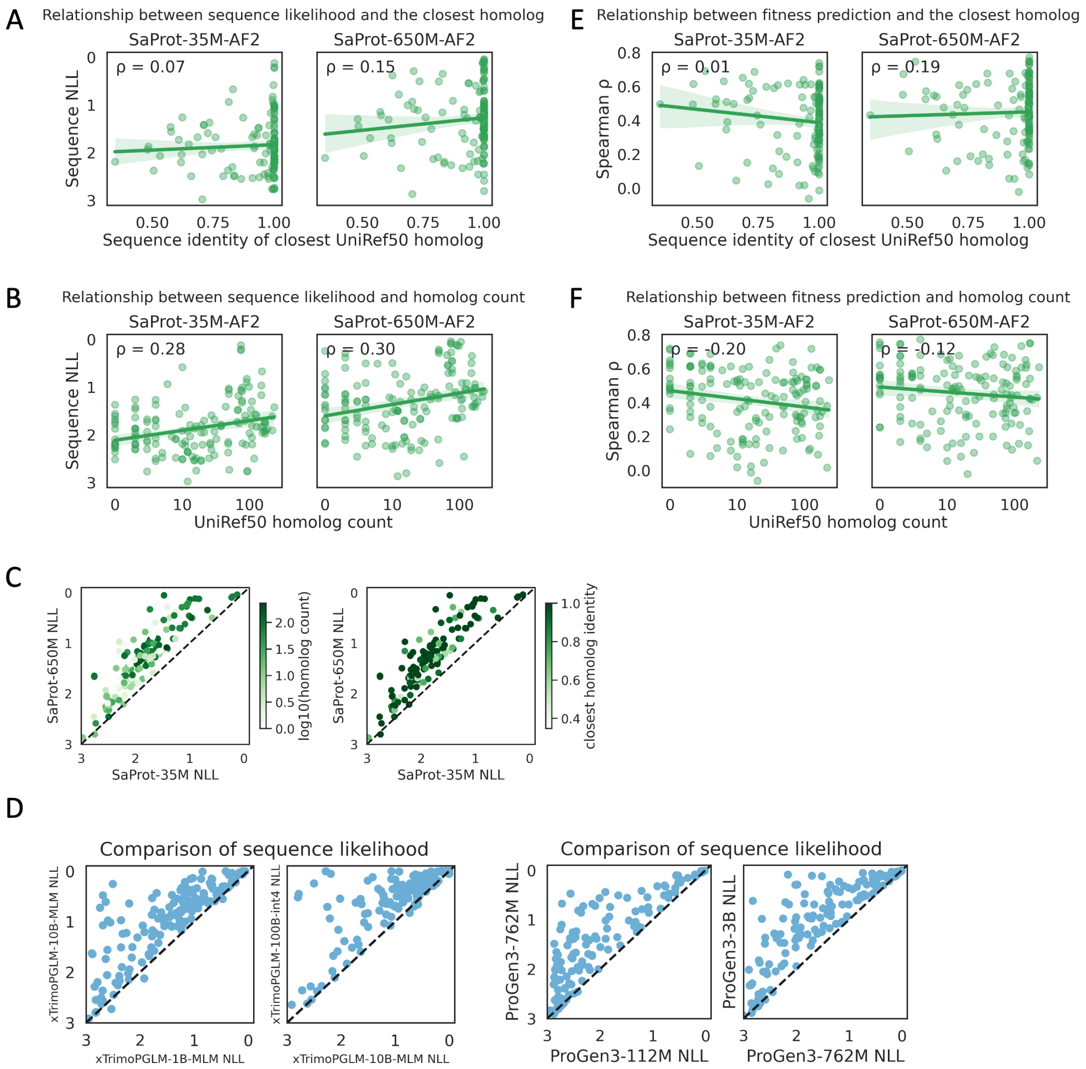


**Figure S2. Analyses of sequence likelihood and fitness prediction performance of other models.**

**A-B**, Relationship between SaProt predicted sequence likelihood and UniRef50 homologs. Each point represents a protein; the x-axes show the number of UniRef50 homologs in the log scale. Homologs are defined as those with ≥20% sequence identity and ≥80% coverage. The curves represent linear regressions, with the shaded areas indicating the 95% confidence intervals. SaProt shares the same training set building procedure as ESM2. **C-D**, Comparison of sequence likelihoods predicted by models of different sizes. **E-F**, Relationship between SaProt fitness prediction performance and UniRef50 homologs.

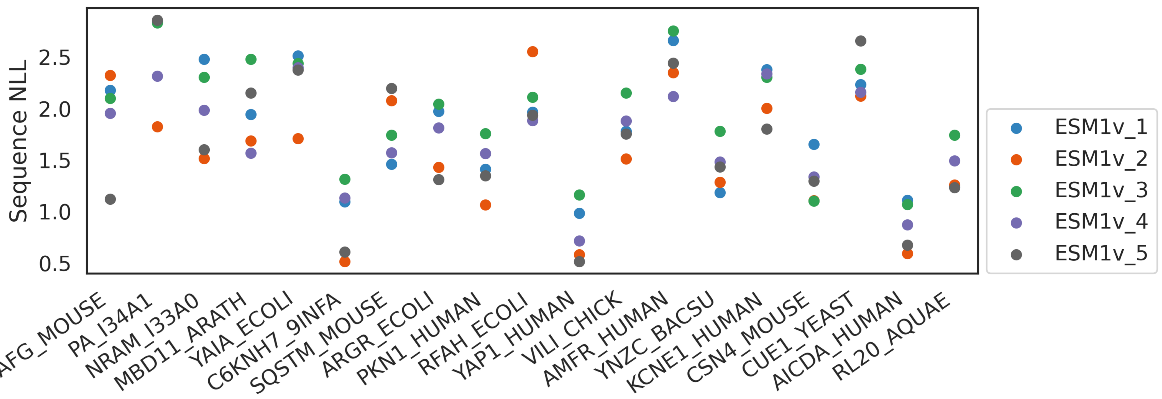


**Figure S3. Sequence likelihood predicted by five ESM1v models.**

All 187 unique wild-type sequences in ProteinGYM (217 experiments, no filtering) are analyzed, only proteins with sequence NLL differences larger than 0.5 among five ESM1v models are shown.

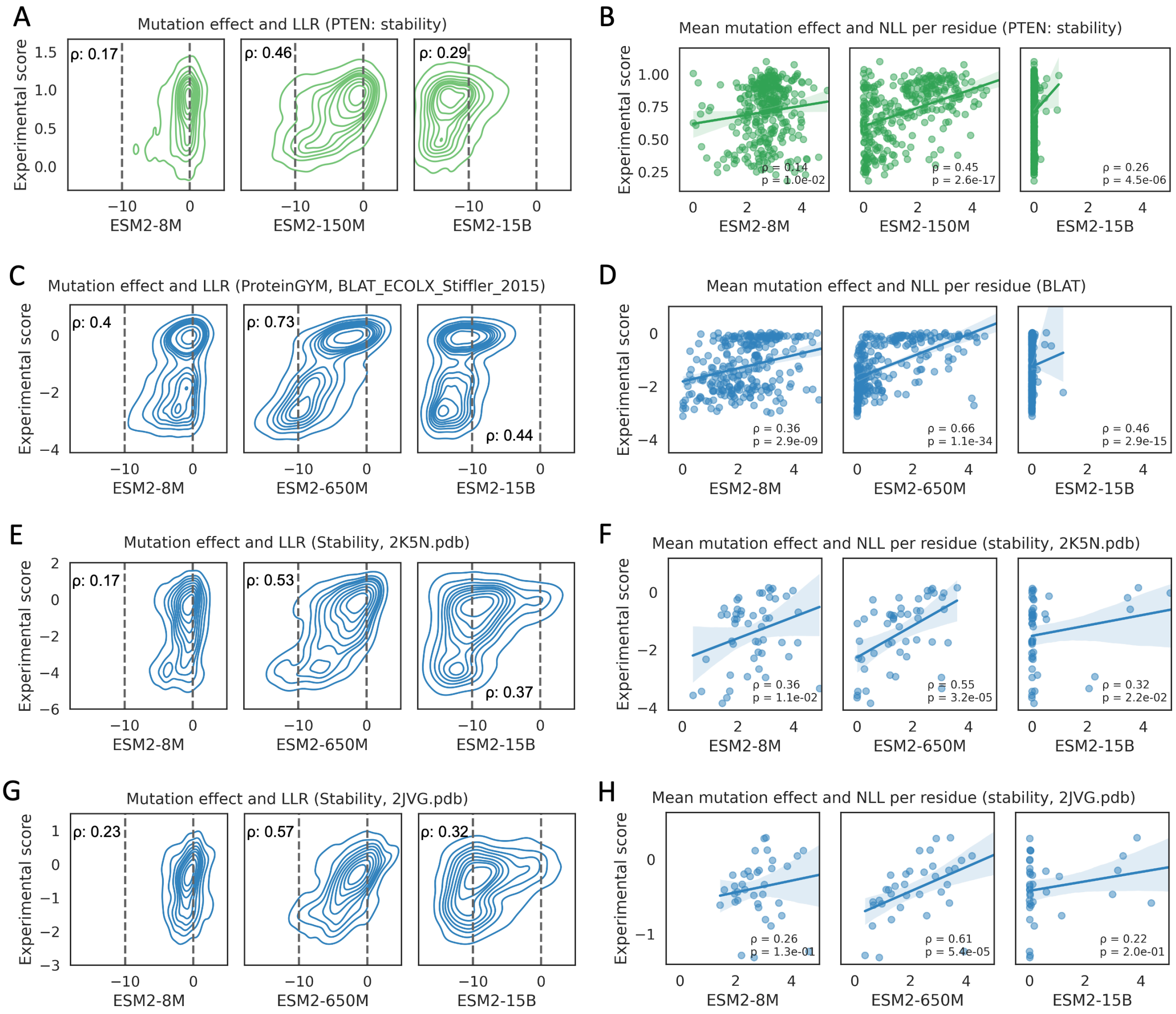


**Figure S4. The relationship between ESM2 prediction and experimental effects.**

**Left:** Relationship between ESM2-predicted LLRs and experimental effects; LLRs of ESM2 are calculated using the masked marginal approach. $\rho$: Spearman correlation. **Right**: Relationship between ESM2-predicted probability per residue (quantified using NLL) and mean experimental effects for mutations at each residue. Each point represents a residue; the lines show the linear regression fit. Spearman correlation coefficients and corresponding p-values are labeled. The visually weaker trends observed in S4F and S4H and relatively larger p-values are due to their relatively small sample sizes, as these proteins are smaller. Experimental scores represent the abundance ratio relative to wild type in an engineered HEK293T cell line in **A–B**, the logarithm of the allele counts in the selected population (ampicillin selection) versus the unselected population, relative to the wild-type in **C–D**, and the ΔΔG of folding stability change in **E–H** from the mega-scaling protein stability dataset.

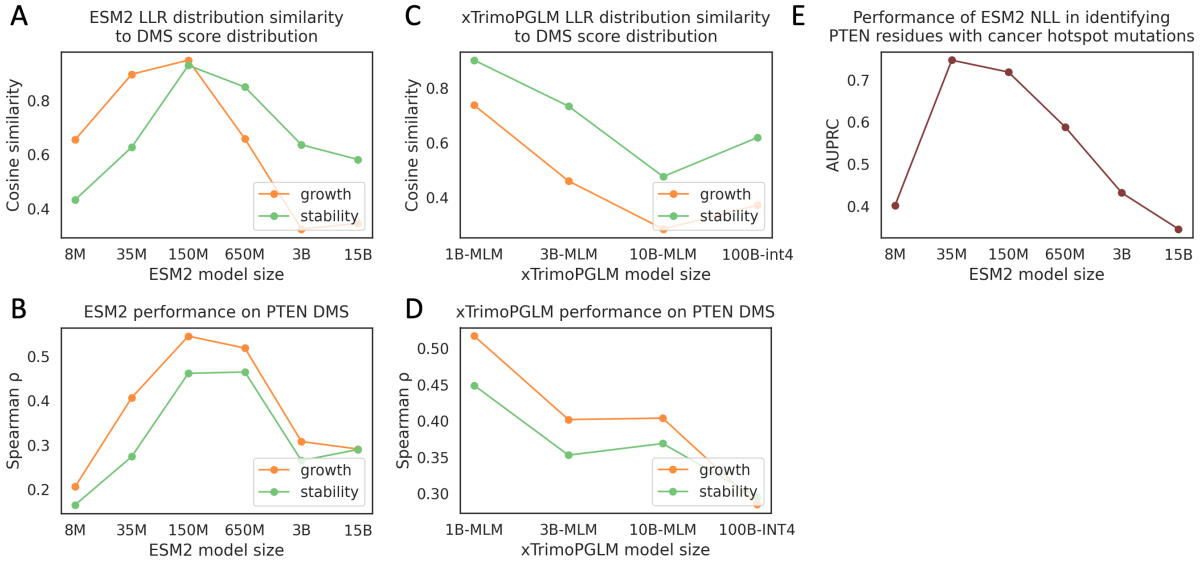


**Figure S5. Analyses of PTEN.**

**A, C**, Similarity between the distributions of model-predicted log-likelihood ratios (LLRs) and experimental mutation effects for two PTEN DMS experiments. To quantify similarity, both predicted and experimental values are discretized into 50 equally spaced bins, distributional similarity is measured using cosine similarity between frequencies across 50 bins. **B, D**, Performance on two PTEN DMS experiments. **E**, Performance of ESM2-predicted residue-level likelihood (NLL) in identifying PTEN residues harboring cancer hotspot mutations (from cBioPortal).

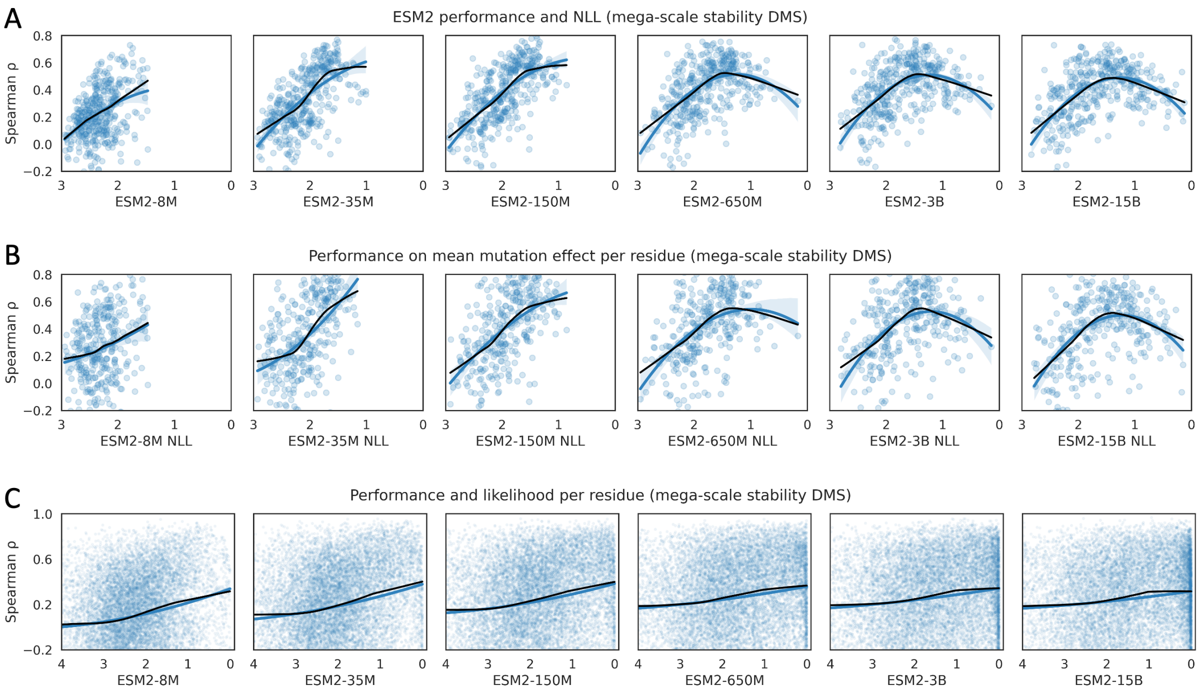


**Figure S6. Relationship between ESM2-predicted likelihood and performance on the mega-scale protein folding stability dataset.**

**A**, The y-axes show the Spearman correlation between LLRs (calculated using the masked marginal approach) and experimental mutation effects, the x-axes represent the negative log-likelihood (NLL) of wild-type sequences. Each point represents an experiment. **B**, The y-axes show the Spearman correlation between mean LLRs and mean experimental mutation effects per residue in each protein, reflecting model understanding of protein context. The x-axes represent the NLL of wild-type sequences. Each point represents an experiment. **C**, The y-axes show the Spearman correlation between LLRs and experimental effects of all 19 mutations per residue, reflecting model understanding of substitution specificity. The x-axes represent the negative log predicted probability of each residue. Each point represents a residue. The black curves show LOWESS modeling, while the blue curves show second-order polynomial regressions, with shaded areas representing the 95% confidence intervals.

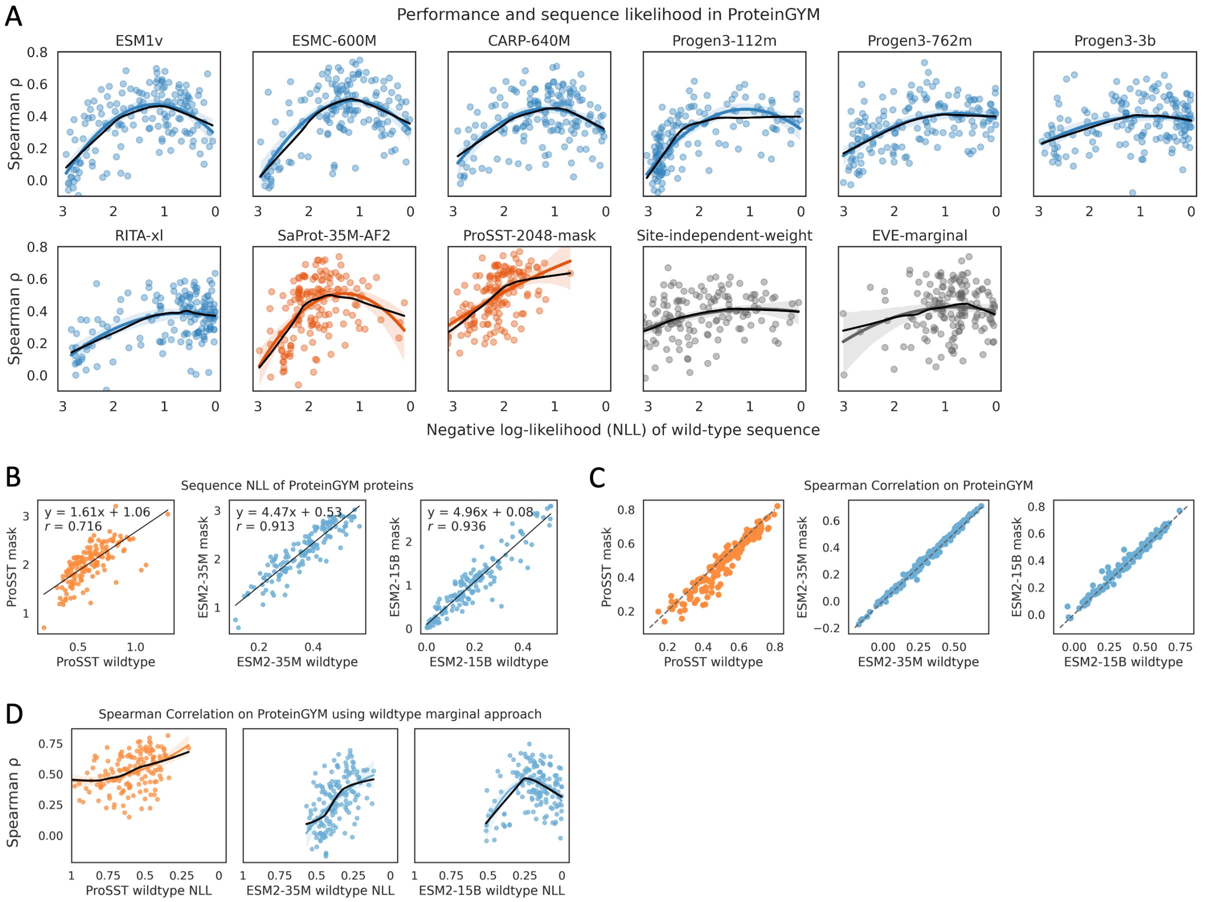


**Figure S7.**

**A**, Relationship between fitness prediction performance and model-predicted sequence likelihood. The y-axes show the Spearman correlation between LLRs and experimental mutation effects, the x-axes represent the negative log-likelihood (NLL) of wild-type sequences. LLRs of ESM models, CARP, SaProt, and ProSST are calculated using the masked marginal approach. LLRs of Progen3 and RITA are calculated using the full sequence. The LLR of the site-independent-weight model is computed as the log difference in amino acid frequency at each site after weighting homologous sequences. See Methods for the EVE-marginal model. Only residues with mutations are included in the NLL calculation. Each point represents an experiment from ProteinGYM (154 experiments after filtering). The black curves show LOWESS modeling, while the colored curves show second-order polynomial regressions, with shaded areas representing the 95% confidence intervals. See Results for the analysis of ProSST, see Discussion for autoregressive pLMs RITA and ProGen3. **B**, For masked language models trained with a fraction of tokens substituted and a fraction of tokens unchanged, likelihoods calculated from the masked marginal and wild-type marginal approaches are linearly correlated. As ProSST is not a pure language model and still has access to the structure token of the masked residue, the slope and correlation differ from those of ESM2. **C**, The log-likelihood ratios (LLRs) computed using these two approaches exhibit nearly identical performance in fitness prediction. **D**, Using the wild-type marginal approach for all methods introduces a systematic shift in the x-axis (i.e., NLL of the wild-type sequence), without changing the overall bell-shaped pattern. As shown, ESM2-15B exhibits a clear bell-shaped trend, consistent with results using the masked marginal approach. Both ProSST and ESM2-35M do not reach the likelihood range where performance declines, and thus only one side of the bell-shaped trend is observed, consistent with the masked marginal approach.

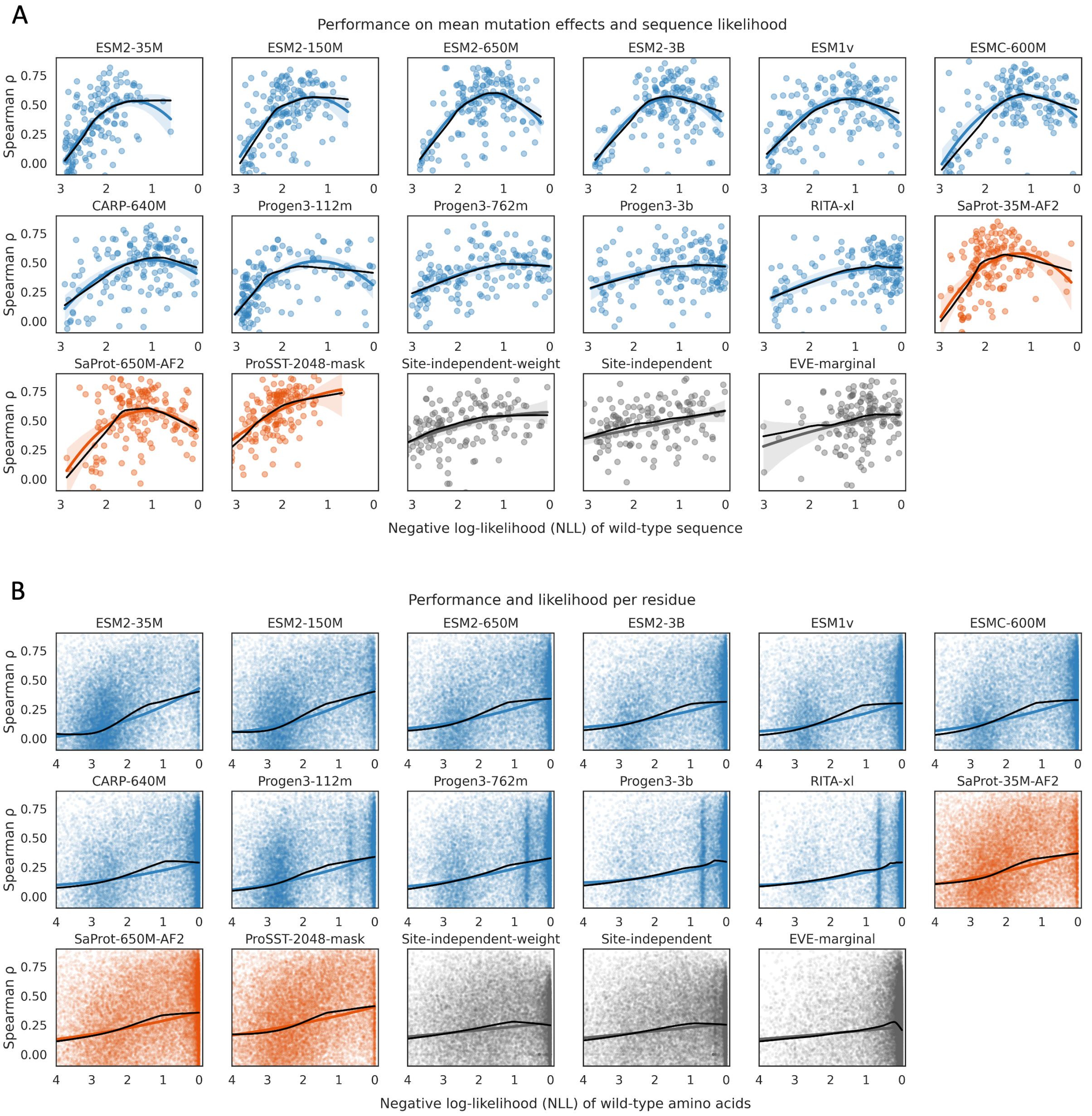


**Figure S8. Model performance on mean mutation effects and mutation effects per residue.**
**A**, The y-axes show the Spearman correlation between mean LLRs and mean experimental mutation effects per residue within each protein, reflecting models' understanding of protein context. The x-axes represent the NLL of wild-type sequences. Only residues with mutations are included in the NLL calculation. Each point represents an experiment. **B**, The y-axes show the Spearman correlation between LLRs and effects of all 19 possible mutations per residue, reflecting substitution specificity. The x-axes represent the negative log predicted probability of each residue. Each point represents a residue. The black curves show LOWESS modeling, while the colored curves show second-order polynomial regressions, with shaded areas representing the 95% confidence intervals.

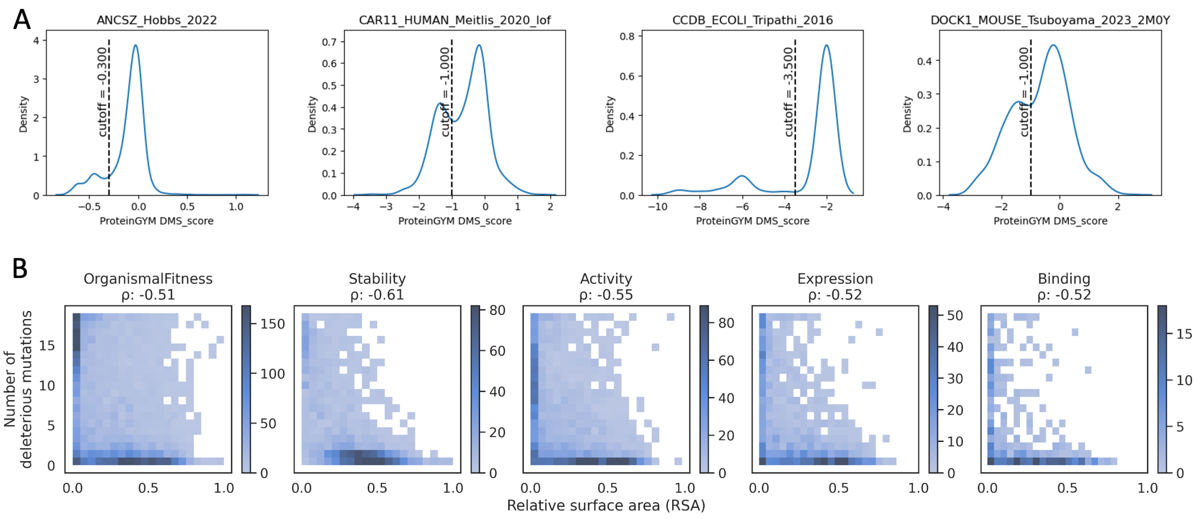


**Figure S9.**

**A**, Example ProteinGYM datasets showing bimodal distributions of mutation effects. Only single-residue substitutions are analyzed. Dashed lines indicate the cutoffs we defined by visual inspection of the bimodal distribution; these cutoff values are provided in Supplementary Data 1 for reproducibility. Refer to ProteinGYM and original publications for the definitions of these DMS_scores. **B**, The relationship between relative surface area (RSA) and number of deleterious mutations per residue. Both RSA and the number of deleterious mutations are discretized into 20 bins, and the color indicates the number of residues within each bin. 122 DMS experiments showing bimodal distributions of mutation effect were analyzed. The pattern observed reflects the relationship between protein structure and mutation sensitivity. The protein core (low RSA) determines structural stability, and thus more residues in the core are sensitive to mutations affecting stability. As other functions, including binding, rely on a stable structure, mutations in the core can also impact these functions. In contrast, mutations at surface residues (high RSA) have relatively small effects on stability on average, with only a fraction involved in specific functions. Because the binding interface is usually small compared to the protein core, it is reasonable that fewer surface residues are mutation-sensitive. One issue is that not many surface residues have mutations that impact binding. We think this might be due to three reasons: the limited number of assays analyzed (eight binding assays analyzed), binding interfaces being buried in AlphaFold predictions, and experiments not being well designed to specifically probe binding.

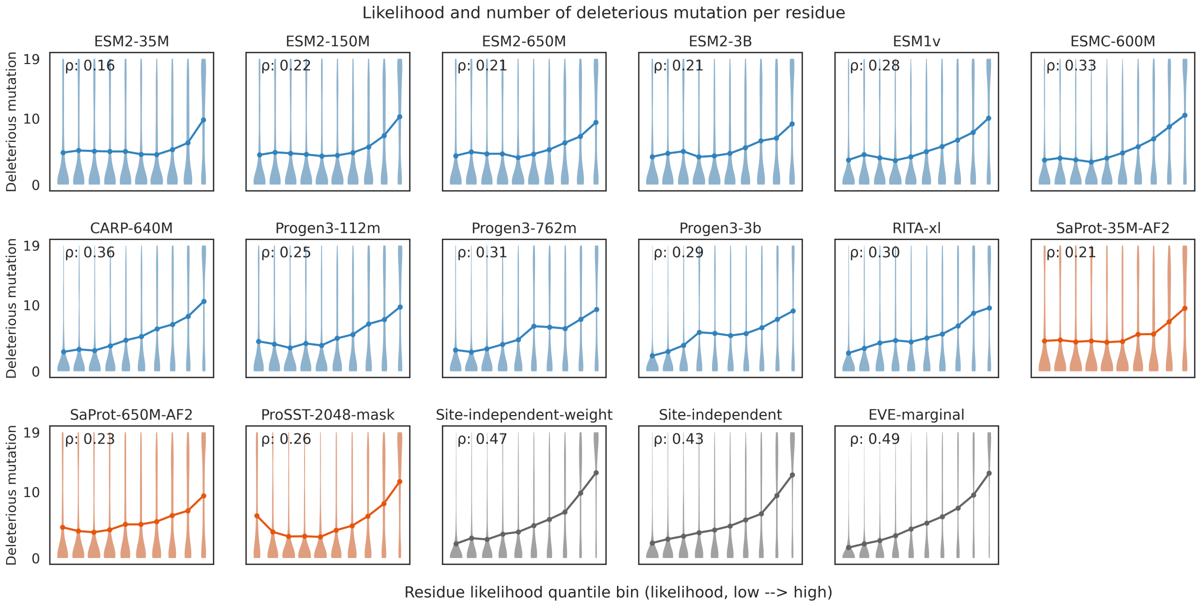


**Figure S10. Comparison of model-predicted residue likelihood with mutation sensitivity.**
Residues are grouped into ten quantile bins based on model-predicted likelihood. For each bin, the distribution and mean value are shown for the number of deleterious mutations per residue (range 0–19).

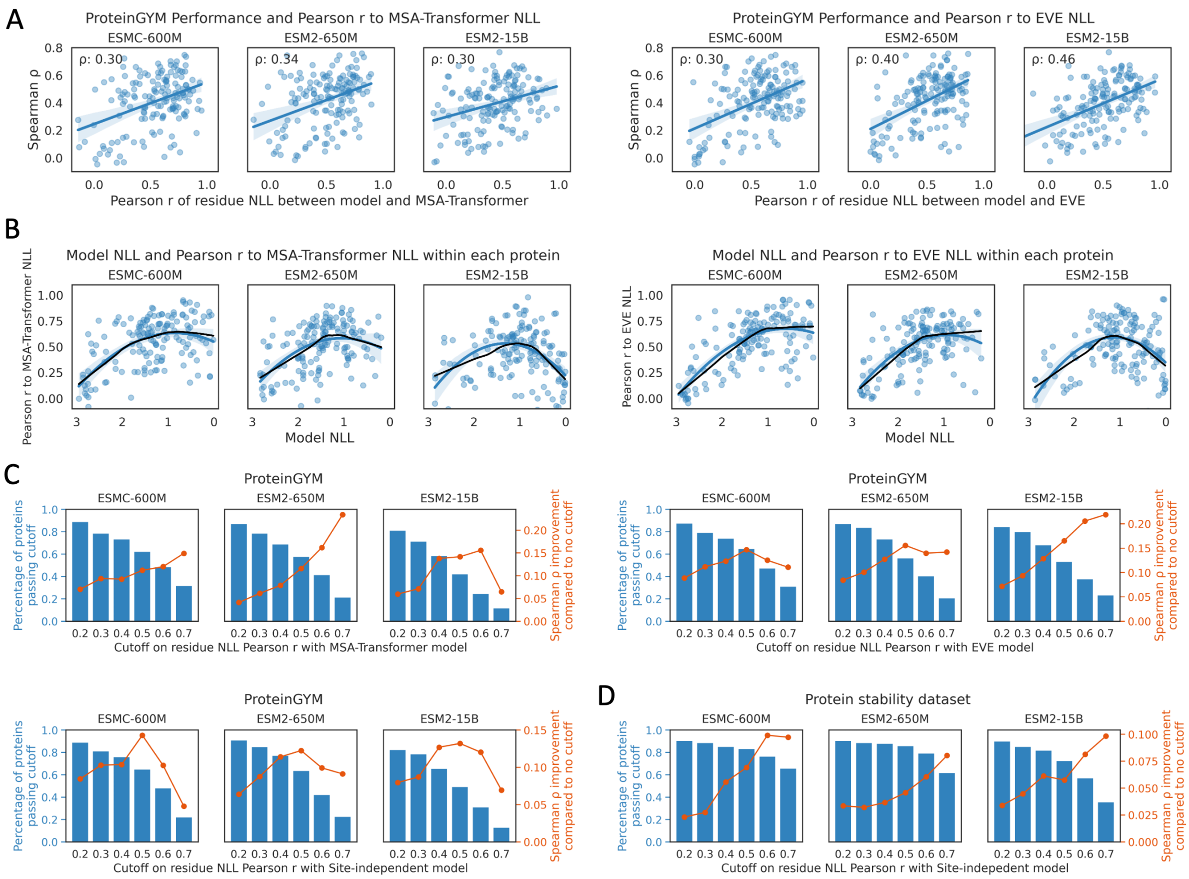


**Figure S11. Comparing ESM models to family-specific models.**

**A**, Relationship between model fitness prediction performance and correlation to NLL from family-specific models. The y-axes show fitness prediction performances of ESM models. The x-axes show the per-residue NLL Pearson correlation between the model and a family-specific model within each protein. Each point represents a ProteinGYM experiment. The curves represent linear regressions, with the shaded areas indicating the 95% confidence intervals. **B**, Relationship between model sequence likelihood and per-residue NLL correlation to family-specific models. The y-axes show the per-residue NLL Pearson correlation between the model and family-specific model within each protein. The x-axes show the model NLL. Each point represents an experiment. The black curves show LOWESS modeling, while the colored curves show second-order polynomial regressions, with shaded areas representing the 95% confidence intervals. **C-D**, Comparison with family-specific models prior to applying general models for fitness prediction. We suggest using the general model only when the model-predicted residue-level NLLs are correlated with those from family-specific models. Considering both the percentage of proteins that pass the correlation cutoff and the performance improvement relative to no filtering (i.e., comparing the mean performance on the selected assays with that across all assays), we recommend applying ESM models when the Pearson correlation with a family-specific model exceeds 0.5. For the mega-scale stability dataset analyzed in panel D, proteins overlapping with ProteinGYM are removed, only protein with number of homologs more than 10 times length are analyzed.

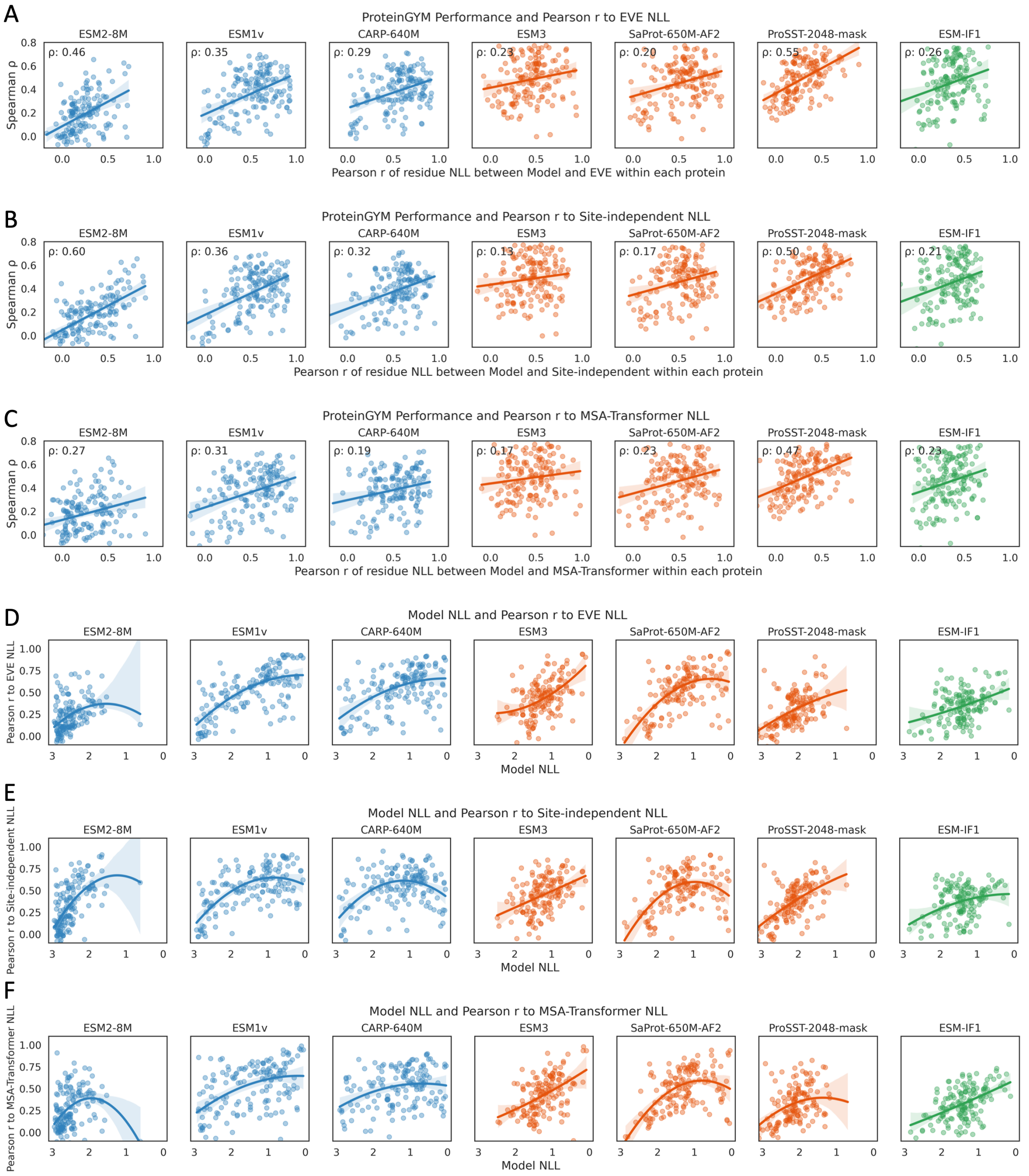


**Figure S12. Relationship between fitness prediction performance and similarity to EVE residue likelihoods.**

**A-C**, Relationship between model fitness prediction performance and per-residue NLL correlation to family-specific models. The y-axes show fitness prediction performance. The x-axes show the per-residue NLL Pearson correlation between the model and a family-specific model within each protein. Each point represents an experiment. The curves represent linear regressions, with the shaded areas indicating the 95% confidence intervals. **D-F**, Relationship between model sequence likelihood and per-residue NLL correlation to family-specific models. The y-axes show the per-residue NLL Pearson correlation between the model and family-specific model within each protein. The x-axes show the model NLL. Each point represents an experiment. The curves show second-order polynomial regressions, with shaded areas representing the 95% confidence intervals.

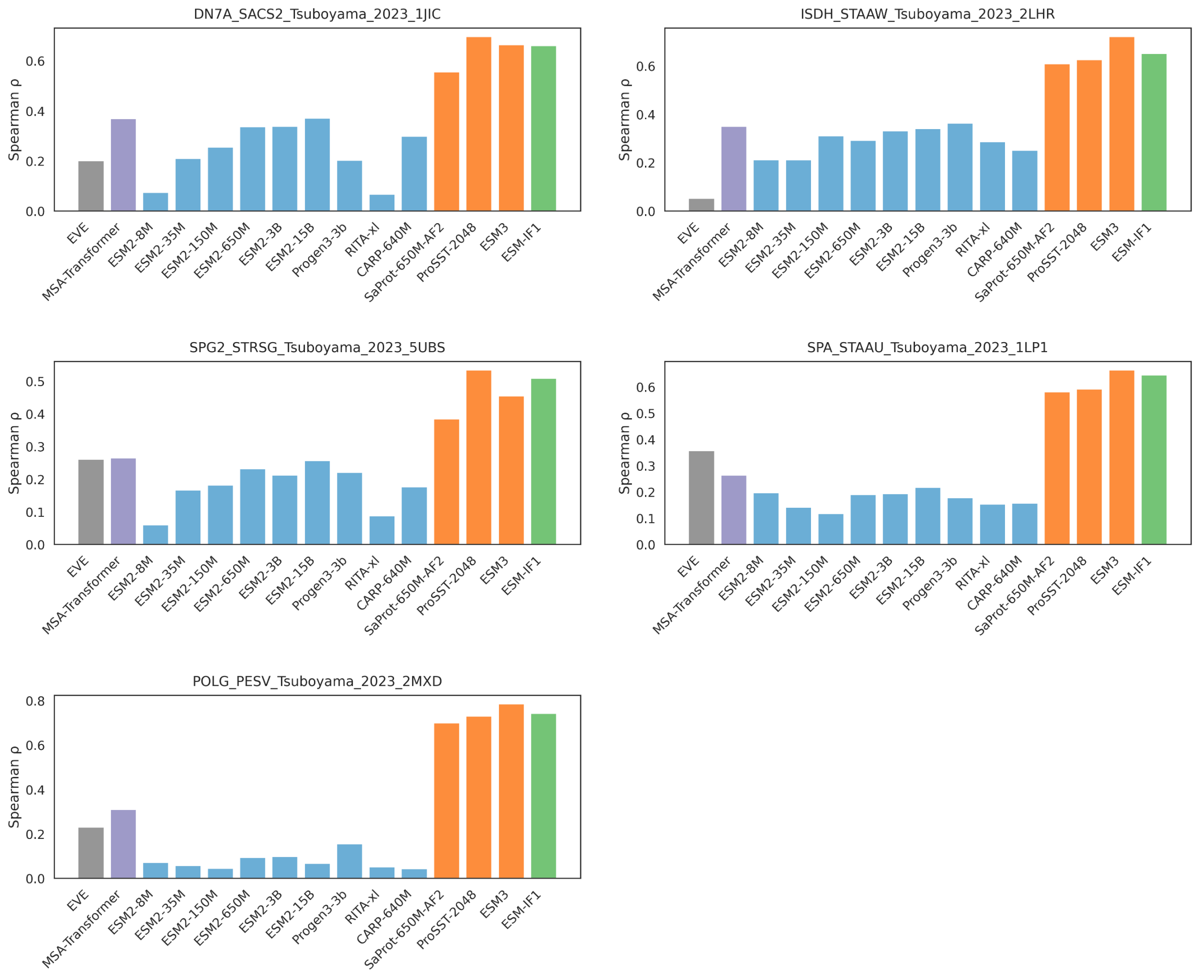


**Figure S13. ProteinGYM assays where structure-informed models outperform sequence-only pLMs and family-specific models.**

All five DMS assays measure mutation effect on stability. Structure-informed models are colored in orange and green.

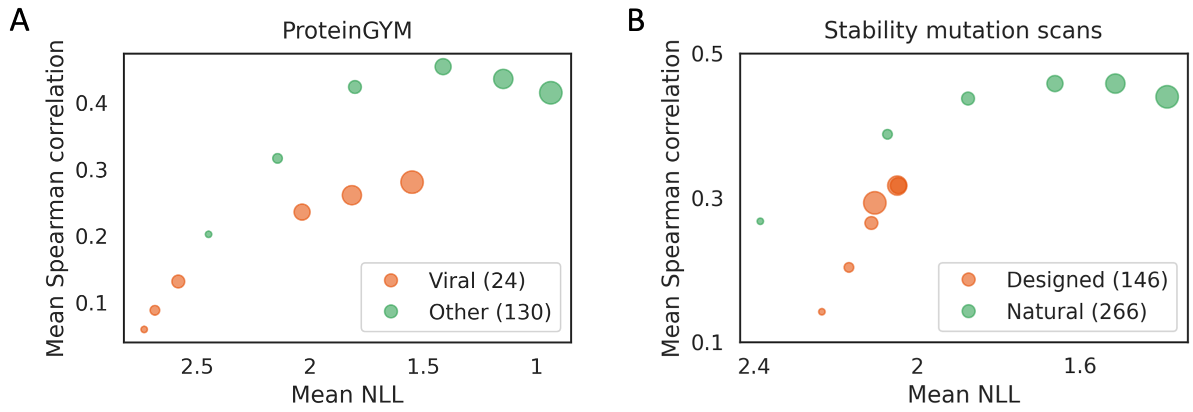


**Figure S14. ESM2 performance on different proteins.**
The mean performance on fitness prediction (average of per-protein Spearman correlation), and predicted sequence likelihood of ESM2 models on DMS experiments in the ProteinGYM benchmark (**A**) and the mega-scale protein stability dataset (**B**). Point size indicates ESM2 model size (from 8M to 15B, same to main Figure 2D).
